# MEF-2 repression of GEM-4/Copine expression decreases muscle excitability by increasing gap junction coupling

**DOI:** 10.64898/2026.04.09.717484

**Authors:** Luna Gao, Edward Pym, Stephen Nurrish, Joshua M. Kaplan

## Abstract

Activity induced gene expression promotes behavioral plasticity (including several forms of learning and memory) by altering synaptic connectivity and neuron excitability. While several transcriptional targets modify synapses, little is known about targets regulating excitability. Here, we identify a single MEF-2 and CRH-1/CREB transcriptional target, *gem-4*/Copine, whose expression increases the excitability of *C. elegans* body muscles. In muscles, *gem-4* expression is repressed by MEF-2 and is induced by CRH-1. GEM-4 expression in muscles decreases the number of gap junctions, thereby increasing membrane resistance and excitability. These results identify activity induced copine expression as a mechanism to control gap junction coupling and excitability.

## Introduction

Activity regulated gene transcription (ART) is mediated by calcium regulated transcription factors (TFs). For example, cAMP response element binding protein (CREB), in association with its co-activators (CBP-1 and CRTC-1), typically induces target gene expression ^1,2^. By contrast, Myocyte enhancer factor 2 (MEF2) can act as either a transcriptional activator or repressor depending on associated co-factors ^3,4^. Transcriptional activation is mediated by c-terminal domains in MEF2 proteins ^5^. MEF2’s transcriptional repressor function is mediated by its association with a co-repressor, class IIa histone deacetylase (e.g. HDAC4). ^6,7^ Increased cytoplasmic calcium activates type 1 and 4 calcium and calmodulin (CaM) dependent protein kinase (CAMK1/4), which activates CREB ^8,9^ and disrupts MEF2-HDAC4 complexes ^10,11^, both leading to increased target expression.

ART is a mechanism for coupling recent activity to future neurodevelopmental and behavioral outcomes ^12,13^. In particular, many results suggest that activity regulated TFs play important roles in memory. CREB has been implicated in long term memory in *Aplysia*, *Drosophila*, and mice ^14–16^. Mef2 inhibits spatial and fear memory in mice ^17^. Expression of a third activity regulated TF (FOS) is used to identify sets of neurons that participate in memory storage and recall ^18^, which are termed engrams, and FOS is required for spatial memory in mice^19^.

In addition to behavioral plasticity, ART has also been proposed to play an important role in in the pathophysiology of neurodevelopmental and psychiatric disorders ^20^. Mutations in CBP (a CREB associated co-activator) are found in Rubinstein-Taybi syndrome, while mutations in both MEF2 and its associated co-repressor HDAC4 are found in Autism spectrum disorder (ASD) ^21–24^. For these reasons, there is significant interest in identifying the mechanisms by which CREB and MEF2 promote behavioral and developmental plasticity.

ART is proposed to modify behavior and neurodevelopment by altering synapse development and function. Mef2 promotes activity induced synapse elimination ^25^ by increasing Arc expression ^26^, which accelerates AMPA receptor endocytosis ^27,28^. CREB is required for the late phase of long-term potentiation (L-LTP) of synaptic transmission in the hippocampus ^29^. CREB also increases neurotransmitter release at *Drosophila* neuromuscular junctions ^30^. Inhibitory synapse formation in the mouse cortex is regulated by CREB induced BDNF expression ^31^ and by Fos induced *Scg2* expression ^32^. Several other ART targets have also been implicated in synaptic plasticity, including Fn14 ^33^, Homer1a ^34^, and Cpg15 ^35^. Collectively, these results strongly support the idea that an important function of ART is modifying synapse development and function, thereby driving behavioral plasticity.

In parallel with synaptic plasticity, other studies suggest that ART also alters how readily neurons are activated by their environmental and synaptic inputs (which is generically referred to as intrinsic excitability)^36^. For example, CREB expression increases excitability of several neuronal cell types ^37–41^; however, it is not known if other activity regulated TFs also modify excitability. Activity-induced increases in excitability could provide a non-synaptic mechanism for learning and memory^42,43^. In contrast to ART effects on synaptic plasticity, little is known about the transcriptional targets that alter intrinsic excitability.

Prior studies on MEF2 function in muscle have focused on its impact on development ^44^. MEF2 promotes expression of TFs that induce the skeletal and cardiac muscle cell fate ^45^. MEF2 loss of function mutations result in profound developmental defects in skeletal and cardiac muscles in mice, flies, and zebrafish ^46–48^. These developmental deficits have made it difficult to study the potential post-developmental functions of MEF2 in muscles. Consequently, little is known about how MEF2 impacts muscle excitability.

These studies raise several important questions. In addition to adjusting the excitability of neurons, does ART also regulate muscle excitability? Conservation of this function in both neurons and muscles would enhance the impact of ART mediated excitability changes on behavioral and developmental plasticity. Do other activity regulated TFs (i.e. beyond CREB) also regulate excitability? What specific ART targets modify excitability? What is the biophysical mechanism for ART regulation of excitability? In principle, ART could adjust excitability by altering passive membrane properties (e.g. membrane resistance or resting membrane potential) or by altering membrane conductances. Identifying specific biophysical mechanisms will provide important insights into how ART modifies behavior and neurodevelopment.

To begin addressing these questions, we analyze how MEF-2 and CRH-1/CREB alter the intrinsic excitability of *C. elegans* body muscles. We exploit the fact that muscle development is not significantly disrupted in *C. elegans mef-2* mutants ^49^, which allows us to readily examine muscle activity in *mef-2* mutants. We show that MEF-2 inhibits muscle excitability and that it does so by inhibiting the CRH-1/CREB induced expression of the *gem-4* gene (which encodes a Copine). GEM-4/Copine expression increases muscle excitability by decreasing gap junction coupling of adjacent muscles, thereby increasing membrane resistance. These results demonstrate that regulating excitability is an ancient function of ART that extends to *C. elegans* muscles.

## Results

### CRH-1 and MEF-2 antagonistically regulate muscle excitability

Prompted by prior studies showing that CREB increases neuron excitability^37–41^, we asked if muscle excitability is altered by MEF-2 and CRH-1/CREB. For the *mef-2* analysis, we used two alleles, a global knockout, *mef-2(gv1)* (a putative null allele), and a muscle specific knockout, hereafter *mef-2(muscle knockout, mKO)*. The *mef-2(mKO)* mutants were constructed by expressing the CRE recombinase in the body muscles of *mef-2*(*nu752* FLEX) mutants, which contain an allele that is inactivated by CRE (as detailed in the methods). CRH-1/CREB’s effects were analyzed using a putative null allele *crh-1(tz2)*.

Muscle excitability was assessed by measuring the rate of action potentials (APs) evoked by a defined depolarizing electrical stimulus. Higher AP firing rates following a depolarizing stimulus indicates increased muscle excitability, whereas decreased excitability would be indicated by lower AP firing rates following the stimulus. The rate of muscle APs evoked by a depolarizing stimulus (a +4 pA current injection) was significantly increased in *mef-2*(*gv1*) (a null allele) and in *mef-2(mKO)* (the muscle specific knockout) (Fig. 1A,D). Increases in AP rate were observed across a range of depolarizing current injections (Fig. 1C), indicating that *mef-2* mutants exhibit increased excitability over a range of stimulus intensities. Because increased AP rates were seen in both global and muscle specific *mef-2* knockouts, these results suggest that MEF-2 acts in body muscles to decrease excitability.

**Figure 1.**
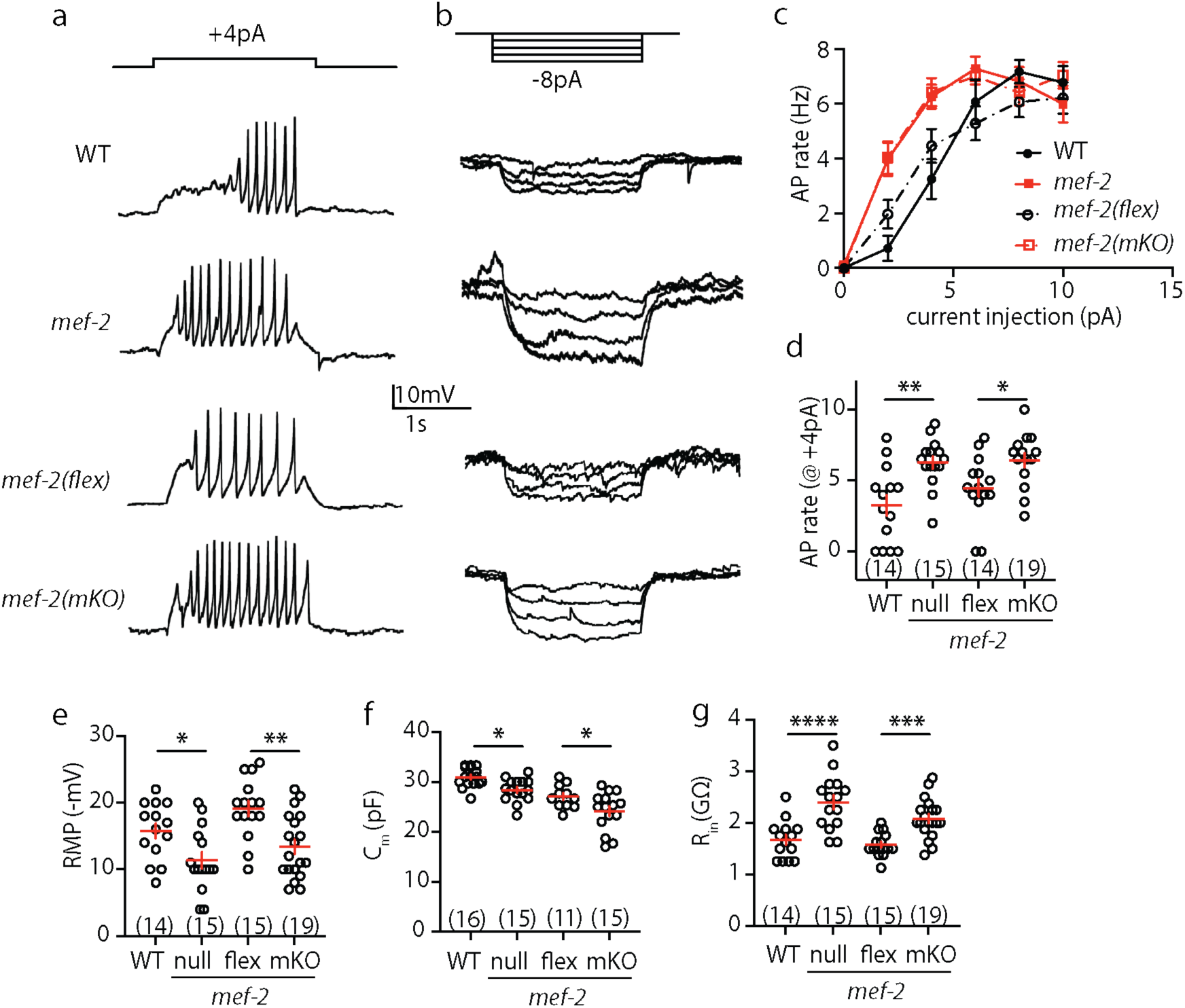
Increased muscle excitability and altered passive membrane properties in *mef-2* mutants. (A-B) Representative traces of APs evoked by depolarizing current (A) and V_m_ changes evoked by hyperpolarizing current (B) are shown in the indicated genotypes. (C) Mean AP rate is plotted as a function of the injected current. (D) Mean AP rate evoked by a +4pA current is plotted. (E-G) Summary data are plotted for mean resting membrane potential (RMP) (E), mean cell capacitance (C_m_) (F), and mean input membrane resistance (R_in_) (G). Two mutations inactivating MEF-2, *mef-2(null)* and *mef-2(mKO)* (a muscle specific knockout) increased the intrinsic excitability of muscles, as indicated by: increase evoked AP rate (A,C,D), depolarized RMP (E), decreased C_m_ (F), and increased R_in_ (B,G). *mef-2(flex)* is the WT control (lacking CRE expression) for *mef-2(mKO)*. Statistical differences were determined by one-way ANOVA with Tukey’s multiple comparisons test. Values that differ significantly are indicated (ns, not significant; *, *p* <0.05; **, *p* <0.01; ***, *p* <0.001; ****, *p* <0.0001). Error bars indicate SEM. Sample sizes for each genotype are indicated in all figure panels.

Because they often oppositely regulate target gene expression, MEF-2 and CRH-1/CREB could have opposite effects on excitability. To test this idea, we analyzed muscle excitability in *crh-1* CREB mutants. Evoked AP rates were not significantly altered in *crh-1* CREB single mutants compared to wild type (WT) controls (Fig. 1, Supplement 1A,B). Thus, inactivating CRH-1/CREB alone was not sufficient to alter excitability, perhaps due to insufficient CRH-1/CREB activation in these growth conditions. If excitability is regulated by genes that are convergently controlled by MEF-2 and CRH-1, the increased excitability observed in *mef-2* mutants could result from increased expression of CRH-1 transcriptional targets. In this scenario, CRH-1 function would be required for the increased excitability observed in *mef-2* mutants. Consistent with this idea, AP firing rates observed in *mef-2(mKO);crh-1* double mutants were not significantly different from those in *crh-1* single mutants (Fig. 2A,C,D). An increased evoked AP rate was restored in *mef-2(mKO);crh-1* double mutants following expression of a *crh-1* cDNA in body muscles (Fig. 2A,C,D). Thus, MEF-2 inactivation increased excitability when CRH-1 is functional (Fig. 1A,C,D) whereas MEF-2 inactivation had no effect on excitability in *crh-1* mutants (Fig. 2A,C,D). These results show that CRH-1 is required for MEF-2’s effects on excitability. Collectively, these results demonstrate that MEF-2 and CRH-1 act in body muscles, where they have opposite effects on excitability.

**Figure 2.**
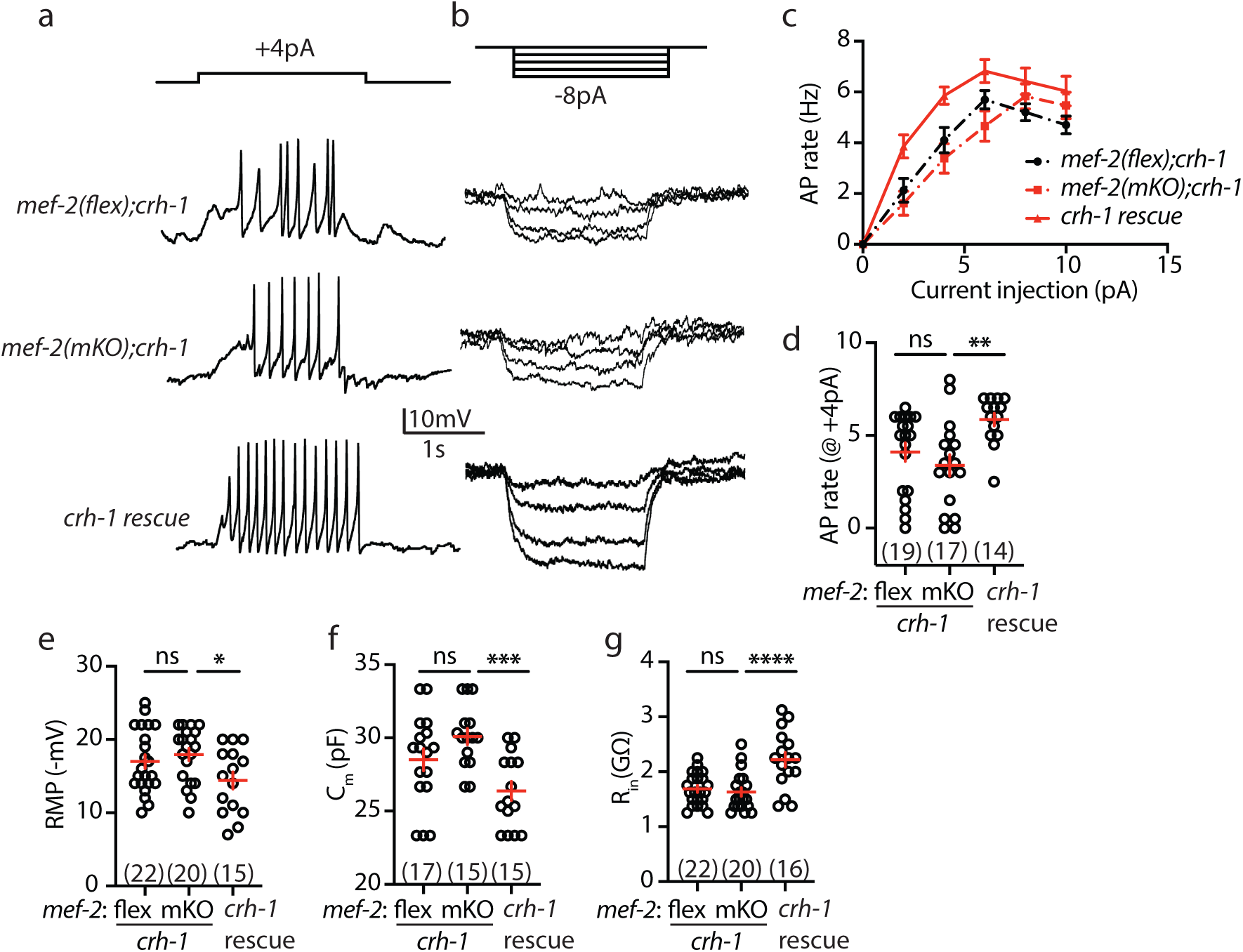
MEF-2’s impact on muscle excitability is eliminated in *crh-1* CREB mutants. (A-B) Representative traces of APs evoked by depolarizing current (A) and V_m_ changes evoked by hyperpolarizing current (B) are shown in the indicated genotypes. (C) Mean AP rate is plotted as a function of the injected current. (D) Mean AP rate evoked by a +4pA current is plotted. (E-G) Summary data are plotted for mean RMP (E), mean C_m_ (F), and mean R_in_ (G). The impact of a *mef-2(mKO)* mutation on evoked AP rate (A,C,D), RMP (E), C_m_ (F), and R_in_ (B,G) was eliminated in double mutants lacking CRH-1 CREB and was restored by a transgene expressing *crh-1* in body muscles (*crh-1* rescue). *mef-2(flex)* is the WT control (lacking CRE expression) for *mef-2(mKO)*. These results suggest that increased muscle excitability in *mef-2* mutants is mediated by increased expression of one or more CRH-1 transcriptional target. Statistical differences were determined as follows: Kruskal-Wallis test with Dunn’s multiple comparisons test (D); one-way ANOVA with Tukey’s multiple comparisons test (E–G). Values that differ significantly are indicated (ns, not significant; *, *p* <0.05; **, *p* <0.01; ***, *p* <0.001; ****, *p* <0.0001). Error bars indicate SEM. Sample sizes for each genotype are indicated in all figure panels.

### Passive membrane properties are altered by *mef-2* and *crh-1* mutations

Increased muscle excitability could arise from a change in passive membrane properties, which refers to electrical characteristics that are not dependent on voltage-activated currents. Passive membrane properties include resting membrane potential (RMP), membrane capacitance (C_m_), and input membrane resistance (R_in_). Increased excitability (as seen in *mef-2* mutants) could result from a depolarized RMP (thereby decreasing the excitatory input required to elicit an AP), or an increased R_in_ (thereby increasing the extent of membrane depolarization following an excitatory stimulus).

To investigate the mechanism for increased excitability, we asked if *mef-2* mutations alter passive membrane properties (Fig. 1B,E,F,G). Consistent with this idea, global [*mef-2(null)*] and muscle specific [*mef-2(mKO)*] *mef-2* knockouts exhibited a slightly depolarized RMP (Fig. 1E), decreased C_m_ (Fig. 1F), and increased R_in_ (Fig. 1B,G). The effects of *mef-2(mKO)* mutations on RMP, C_m_, and R_in_ were eliminated in *mef-2(mKO);crh-1* double mutants and were restored by a transgene expressing CRH-1 in body muscles (Fig. 2B,E,F,G). By contrast, passive membrane properties were unaltered in *crh-1* single mutants (Fig. 1, Supplement 1C-E). These results suggest that MEF-2 and CRH-1/CREB antagonistically control muscle excitability by having opposite effects on passive membrane properties.

### Analysis of voltage-activated currents in *mef-2* mutants

Increased excitability could also result from changes in voltage activated currents. *C. elegans* body muscles have voltage-activated calcium and potassium currents. The voltage-activated calcium currents are mediated by EGL-19/CaV1 channels ^50,51^. Neither the peak amplitude of muscle EGL-19/CaV1 current (Fig. 2, Supplement 1A,B) nor the voltage-dependence of its activation (Fig. 2, Supplement 1C) was significantly altered in *mef-2(null)* mutants. Voltage-activated potassium currents in body muscles are mediated by SHK-1/KCNA and SLO-2/BK channels ^52,53^. SHK-1 channel function can be assessed in recordings using an internal solution containing low chloride levels ^54^. SHK-1 current was modestly reduced in both *mef-2(null)* (Fig. 2, Supplement 1D-F) and *mef-2(mKO)* (Fig. 2, Supplement 2A,B) mutants. SHK-1 current was also reduced in *crh-1* CREB mutants but was not further reduced in *mef-2;crh-1* double mutants (Fig. 2, Supplement 1G-I). The lack of additive effects of *mef-2* and *crh-1* mutations on SHK-1 current suggests that CRH-1 is required for MEF-2’s effects on SHK-1 currents. SLO-2/BK channel function can be assayed in recordings utilizing internal solutions with high chloride levels, which activates SLO-2/BK channels ^55^. To isolate the SLO-2 current, these recordings were done in *shk-1* mutants. SLO-2 current was unaltered in *shk-1;mef-2(null)* double mutants (Fig. 2, Supplement 1J-L). Taken together, these results suggest that MEF-2 and CRH-1 enhance SHK-1 currents but have no effect on SLO-2/BK or EGL-19/CaV1 currents.

### MEF-2 effects on muscle excitability are mediated by changes in gap junction coupling

Thus far, our results suggest that many aspects of intrinsic muscle activity are modified by MEF-2 and CRH-1/CREB. We next did several experiments to determine the mechanism for the altered passive membrane properties in *mef-2* mutants. Membrane capacitance scales with the total area of the plasma membrane available to store charge separation following a shift in membrane potential. Membrane resistance is inversely proportional to the total ion conductance across the plasma membrane, which is mediated by ion channels. Thus, changes in SHK-1 current could contribute to the altered passive membrane properties in *mef-2* mutants. Arguing against this idea, R_in_ and C_m_ were not altered in *shk-1* mutants (Fig. 2, Supplement 2C,D,F) while RMP was hyperpolarized (Fig. 2, Supplement 2E), opposite to the effect seen in *mef-2* mutants (Fig. 1E). The hyperpolarized RMP in *shk-1* mutants was surprising because inactivating a potassium channel would be expected to depolarize RMP. This unexpected RMP hyperpolarization could result from a homeostatic change in other ion channels compensating for the loss of potassium current in *shk-1* mutants ^56^.

An alternative model to explain the observed changes in passive membrane properties is that MEF-2 and CRH-1 regulate gap junctions. Gap junctions mediate direct ion flow between coupled cells, thereby simultaneously impacting membrane resistance and capacitance. Thus, decreased gap junction coupling could potentially account for the increased R_in_ and decreased C_m_ observed in *mef-2* mutants. *C. elegans* body muscles are extensively connected by gap junctions ^57^. As in other invertebrates, *C. elegans* gap junctions are formed by innexin proteins ^58^. To confirm that gap junctions have the expected effects on muscle excitability and passive membrane properties, we analyzed *unc-9* mutants, which lack an innexin that forms body muscle gap junctions ^59^. Here again, we used two alleles for this analysis, *unc-9(nu832)* (a null allele) and *unc-9(nu859* Flox) (a CRE inactivated allele). Muscle specific knockouts, hereafter *unc-9*(*mKO*), were constructed by expressing CRE in body muscles of *unc-9(nu859* Flox) animals. We found that *unc-9* null and *unc-9*(*mKO*) mutants both exhibited increased evoked AP frequency (Fig. 3A-B; Fig. 3, Supplement 1A-B), decreased C_m_ (Fig. 3F; Fig. 3, Supplement 1D), and increased R_in_ (Fig. 3G, Fig. 3, Supplement 1E). These results confirm that decreased gap junction coupling provides a mechanism to increase muscle excitability. RMP was not significantly altered in *unc-9* mutants (Fig. 3E, Fig. 3, Supplement 1C), which is not surprising because RMP is set by ion flow between the cytoplasm and the extra-cellular bath (which is not directly impacted by gap junctions).

**Figure 3.**
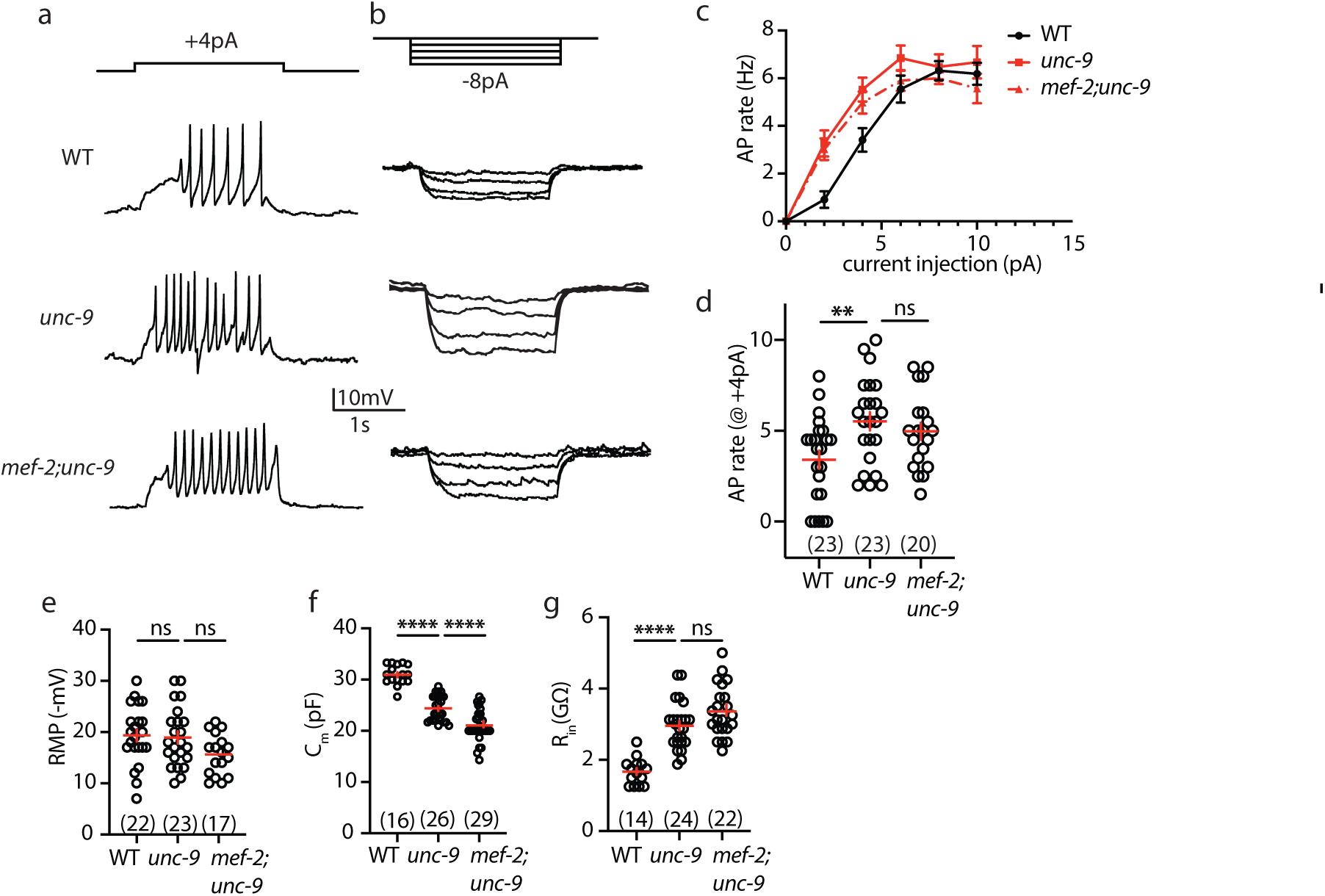
Gap junctions are required for the increased muscle excitability in *mef-2* mutants. (A-B) Representative traces of APs evoked by depolarizing current (A) and V_m_ changes evoked by hyperpolarizing current (B) are shown in the indicated genotypes. (C) Mean AP rate is plotted as a function of the injected current. (D) Mean AP rate evoked by a +4 pA current is plotted. (E-G) Summary data are plotted for mean resting membrane potential (RMP) (E), mean cell capacitance (C_m_) (F), and mean input resistance (R_in_) (G). A mutation inactivating *unc-9* Innexin increases intrinsic muscle excitability, as indicated by increased evoked AP rate (B-D), decreased C_m_ (F), and increased R_in_ (G). In *mef-2;unc-9* double mutants, MEF-2’s impact on evoked AP rate (B-D), RMP (E), and R_in_ (G) was eliminated, while additive effects were seen on C_m_ (F). These results suggest that changes in UNC-9 gap junctions are required for many of MEF-2’s effects on excitability and passive membrane properties. Statistical differences were determined by one-way ANOVA with Tukey’s multiple comparisons test. Values that differ significantly are indicated (ns, not significant; **, *p* <0.01; ****, *p* <0.0001). Error bars indicate SEM. Sample sizes for each genotype are indicated in all figure panels.

Thus far, our results show that *mef-2* and *unc-9* single mutants exhibit similar changes in passive membrane properties (decreased C_m_ and increased R_in_), and similar increases in excitability (increased AP rate). MEF-2 and UNC-9 could function independently to alter excitability by distinct mechanisms. In this scenario, *mef-2* and *unc-9* mutations would have additive effects on excitability in double mutants. Alternatively, MEF-2 could alter excitability by regulating gap junction coupling. In this scenario, *mef-2* and *unc-9* mutations would not have additive effects on excitability in double mutants. To distinguish between these possibilities, we compared the excitability and passive membrane properties of *unc-9* single mutants to those in *mef-2;unc-9* double mutants. We found that evoked AP rate (Fig. 3A,C,D), RMP (Fig. 3E), and R_in_ (Fig. 3B,G) in *unc-9* single mutants were not significantly different from those in *mef-2;unc-9* double mutants. The lack of additive effects of *mef-2* and *unc-9* mutations on AP rate, RMP, and R_in_ suggest that changes in gap junction coupling are required for MEF-2’s effects on excitability. By contrast, additive effects on C_m_ were observed in *mef-2;unc-9* double mutants (Fig. 3F), which could reflect MEF-2 effects on additional innexins (i.e. beyond UNC-9) that contribute to body muscle gap junctions^60^.

To confirm that MEF-2 regulates gap junctions, we used paired recordings of adjacent muscles (Fig. 4A) to assess gap junction coupling, as previously described ^59^. In these recordings, a hyperpolarizing current is injected into one muscle (designated cell 1) and the resulting membrane voltage change (ΔV_m_) is recorded in both cells (i.e. cell 1 and 2) (Fig. 4B-D). Electrical coupling is quantified by measuring the coupling coefficient (which is calculated as the cell 2/cell 1 ΔV_m_ ratio) (Fig. 4E). Using this strategy, we observed strong coupling between adjacent muscles in WT animals and this coupling was significantly reduced in *unc-9* mutants (Fig. 4B-E), as seen in prior studies ^59,60^. Coupling was also significantly reduced in *mef-2(null)* mutants (Fig. 4B-E), consistent with decreased gap junction function. Taken together, these results suggest that increased muscle excitability in *mef-2* mutants is a consequence of decreased gap junction coupling.

**Figure 4.**
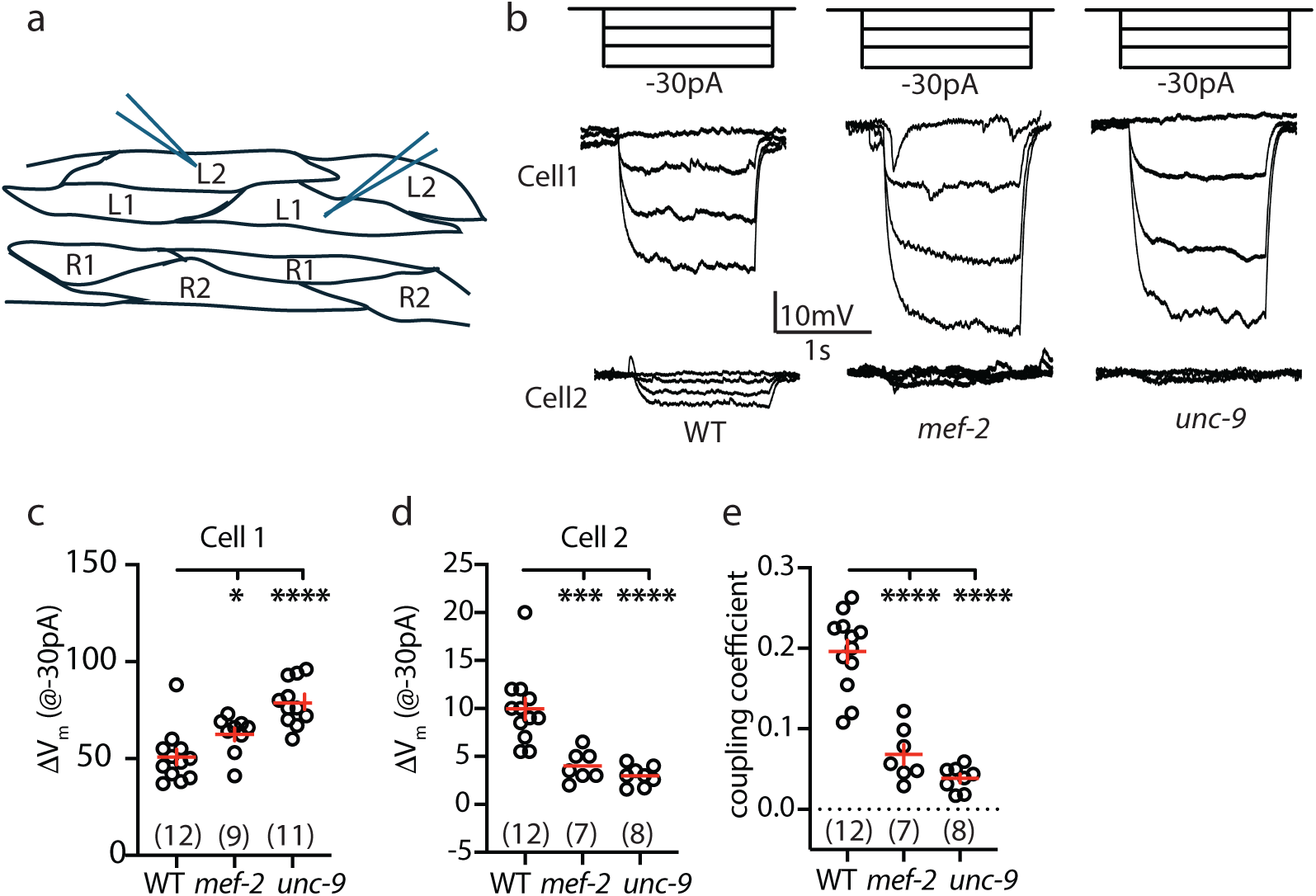
Gap junction coupling is decreased in *mef-2* mutants. Gap junction coupling is assayed in the indicated genotypes by paired recordings of adjacent muscle cells (as illustrated in A). In these recordings (B), hyperpolarizing current is injected into cell 1 and resulting changes in membrane potential (ΔV_m_) are measured in cell 1 and 2. Representative traces (B) and mean ΔV_m_ in cell 1 (C) and cell 2 (D) are shown. Mean gap junction coupling coefficient (E), which is calculated as the cell 2/cell 1 ΔV_m_ ratio, is significantly reduced in *mef-2* and *unc-9* mutants. Statistical differences were determined as follows: one-way ANOVA with Tukey’s multiple comparisons test (E); Kruskal-Wallis test with Dunn’s multiple comparisons test (J, K). Values that differ significantly are indicated (ns, not significant; *, *p* <0.05; ***, *p* <0.001; ****, *p* <0.0001). Error bars indicate SEM. Sample sizes for each genotype are indicated in all figure panels.

### MEF-2 represses baseline *gem-4* expression

Which MEF-2 and CRH-1/CREB transcriptional targets regulate gap junctions? Muscle depolarization induces expression of a broad set of genes via activation of CRH-1/CREB ^1^. The most highly CRH-1/CREB induced gene in depolarized muscles was *gem-4*, which encodes a calcium-binding protein copine (CPNE). Prior studies suggest the CPNE proteins are potentially involved in synaptic plasticity. Mouse CPNE6 expression in hippocampal brain slices is induced by chemical depolarization and by LTP ^61^, indicating that activity induced CPNE expression is conserved across phylogeny. Second, mouse CPNE6 is required hippocampal learning and memory, hippocampal LTP, and LTP induced increases in spine size and surface abundance of synaptic AMPA-type glutamate receptors (GluAs)^62,63^. For these reasons, we decided to focus on *gem-4* CPNE as a candidate downstream effector of MEF-2 and CRH-1/CREB in muscles.

If *gem-4* is a downstream effector for MEF-2’s effects on excitability, *gem-4* expression should be convergently controlled by MEF-2 repression and CRH-1 activation. The *gem-4* promoter contains several predicted MEF2 binding sites and is bound to MEF-2 in chromatin-immunoprecipitation experiments ^64,65^, suggesting that MEF2 could regulate *gem-4* expression (Fig. 5A). To test this idea, we analyzed *gem-4* expression in *mef-2* mutants. For this analysis, we used CRISPR to convert the endogenous *gem-4* locus into an operon containing an upstream cistron expressing histone H2B tagged with the fluorescent protein monomeric StayGold (mSG) (Fig. 5B). Using the *gem-4(nu858* mSG::H2B) allele, we found that mSG fluorescence in body muscle nuclei was significantly increased in *mef-2* null and *mef-2(mKO)* mutants, and this effect was eliminated in *mef-2; crh-1* double mutants (Fig. 5C-D). These results strongly support the idea that *gem-4* expression is activated by CRH-1/CREB and is repressed by MEF-2 (Fig. 5A).

**Figure 5.**
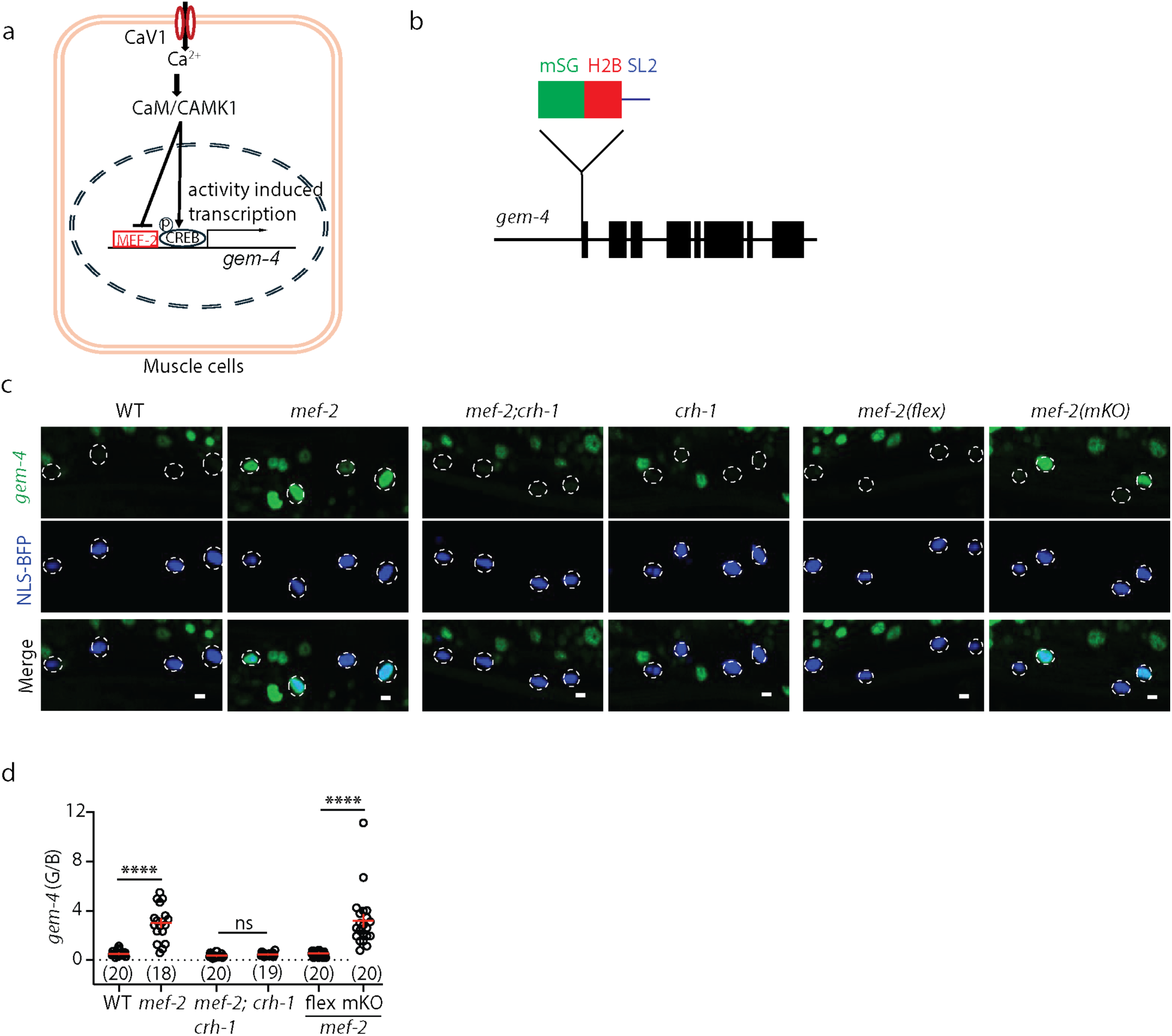
MEF-2 and CRH-1 antagonistically regulate *gem-4* expression in body muscles. (A) Depolarizing muscle induces *gem-4* expression via activation of CRH-1/CREB ^1^. MEF-2 binds the *gem-4* transcription control region in chromatin immunoprecipitation experiments, suggesting that it could also regulate *gem-4* expression ^64,65^. (B) A *gem-4* transcriptional reporter was made by inserting an upstream cistron expressing monomeric StayGold (mSG) tagged H2B (followed by an SL2 splice acceptor) upstream of the endogenous *gem-4* locus. (C-D) Muscle expression of *gem-4* was significantly increased in *mef-2(null)* and *mef-2(muscle knockout, mKO*) mutants and this effect was blocked in *mef-2; crh-1* double mutants. *mef-2(flex)* indicates the parental strain for *mef-2(mKO)*, which lacks the CRE expressing transgene. Body muscle nuclei (dashed circles) were identified by a transgene expressing nuclear localized BFP (NLS-BFP) with the *myo-3* promoter. Expression of *gem-4(nu858* mSG::H2B) was quantified as the mSG/BFP (G/B) fluorescence ratio in muscle nuclei. Representative images (C) and mean *gem-4* expression (D) are shown in the indicated genotypes. Statistical differences were determined by one-way ANOVA with Tukey’s multiple comparisons test. Values that differ significantly are indicated (ns, not significant; ****, *p* <0.0001). Error bars indicate SEM. Sample sizes for each genotype are indicated in all figure panels. Scale bar indicates 4 μm.

### GEM-4/CPNE is required for MEF-2 effects on gap junction coupling

Next, we asked if *gem-4* CPNE contributes to changes in muscle excitability in *mef-2* mutants. Mutations inactivating *gem-4* CPNE eliminated the effect of *mef-2(null)* mutations on evoked AP frequency (Fig. 6A,C,D), RMP (Fig. 6E), C_m_ (Fig. 6F), and R_in_ (Fig. 6G). These results 12 indicate that GEM-4 CPNE is required for *mef-2* mutant defects in excitability and passive membrane properties. To confirm that GEM-4 increases excitability by decreasing gap junction coupling, we used paired cell recordings to directly measure electrical coupling of adjacent muscles. The gap junction coupling coefficient observed in *gem-4* single mutants was not significantly different from that in *mef-2;gem-4* double mutants (Fig. 7A-D), indicating that inactivating *gem-4* eliminates the effects of *mef-2* mutations on gap junction coupling. By contrast, *gem-4* mutations had no effect on SHK-1 currents in *mef-2* mutants (Fig. 6, Supplement 1A-B). These results suggest that GEM-4/CPNE is required for many (but not all) of MEF-2’s effects on muscle excitability.

**Figure 6.**
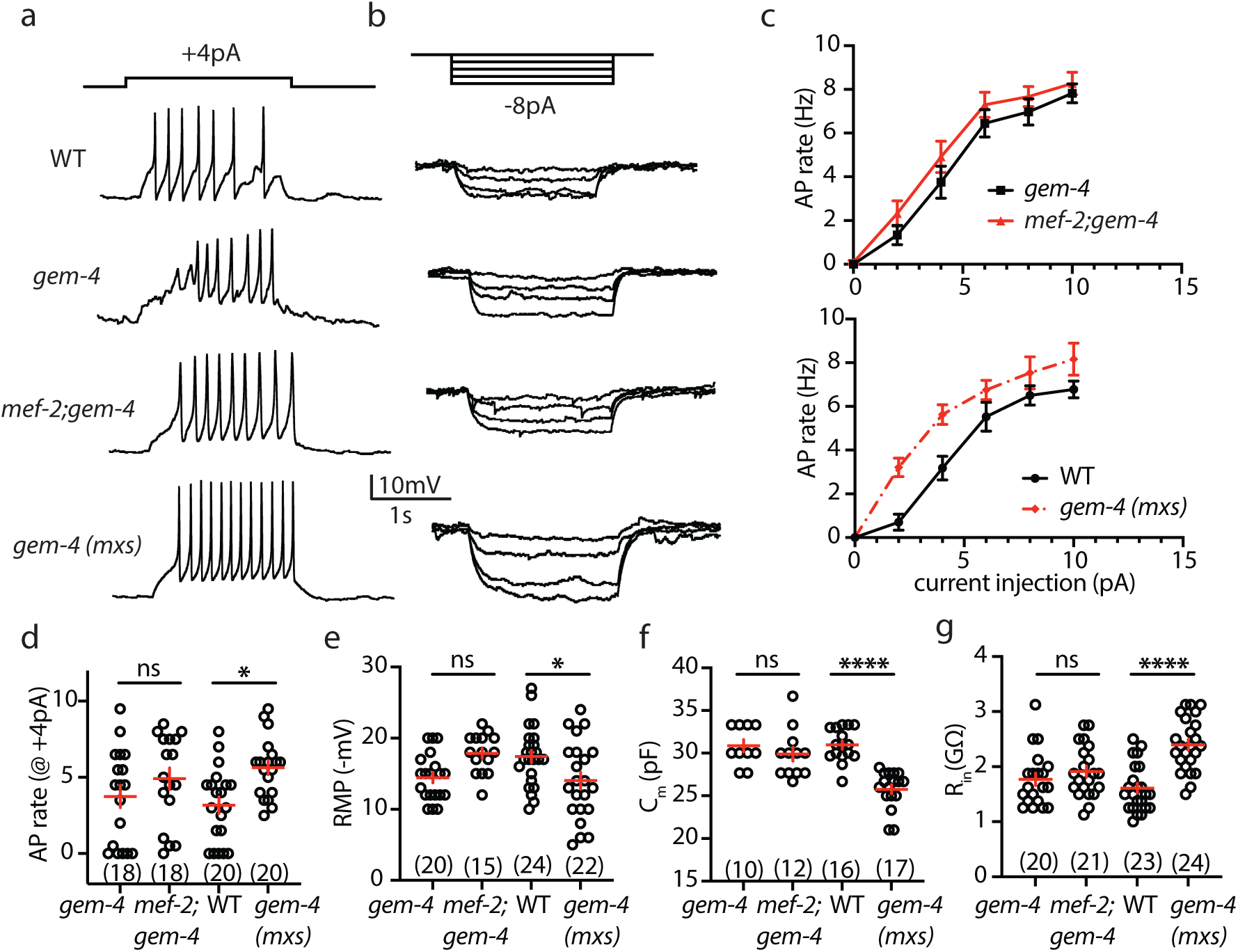
GEM-4 CPNE mediates MEF-2’s effects on excitability and passive membrane properties. (A-B) Representative traces of APs evoked by depolarizing current (A) and V_m_ changes evoked by hyperpolarizing current (B) are shown in the indicated genotypes. (C) Mean AP rate is plotted as a function of the injected current. (D) Mean AP rate evoked by a +4 pA current is plotted. (E-G) Summary data are plotted for mean resting membrane potential (RMP) (E), mean cell capacitance (C_m_) (F), and mean input resistance (R_in_) (G). The impact of a *mef-2* null mutation on intrinsic muscle excitability and passive membrane properties is eliminated in *mef-2;gem-4* double mutants. By contrast, a transgene constitutively expressing GEM-4 in body muscles, *gem-4(mxs)*, mimics the *mef-2* mutant phenotypes of increased excitability and altered passive membrane properties. These results suggest increased and decreased GEM-4 CPNE function have opposite effects on excitability and passive membrane properties. Statistical differences were determined by one-way ANOVA with Tukey’s multiple comparisons test. Values that differ significantly are indicated (ns, not significant; *, *p* <0.05; ****, *p* <0.0001). Error bars indicate SEM. Sample sizes for each genotype are indicated in all figure panels.

**Figure 7.**
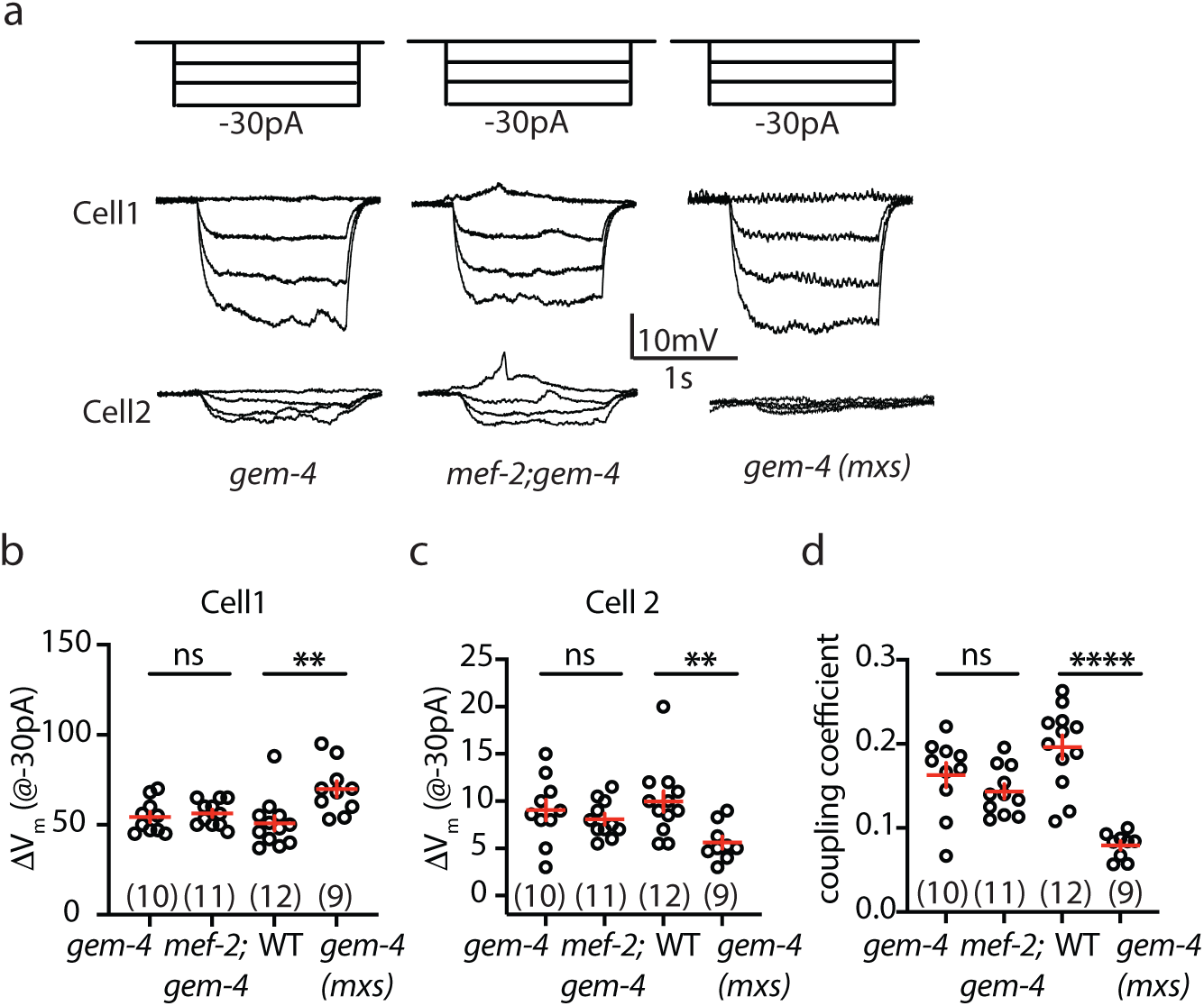
GEM-4 CPNE decreases gap junction coupling. Gap junction coupling is assayed in the indicated genotypes by paired recordings of adjacent muscle cells. In these recordings (A), a hyperpolarizing current is injected into cell 1 and resulting changes in membrane potential (ΔV_m_) are measured in cell 1 and 2. Representative traces (A), mean ΔV_m_ in cell 1 (B) and cell 2 (C), and mean coupling coefficients (D) are shown. The impact of a *mef-2* null mutation on gap junction coupling is eliminated in *mef-2;gem-4* double mutants. By contrast, a transgene over-expressing GEM-4 in body muscles, *gem-4(mxs)*, decreases gap junction coupling. These results suggest increased and decreased GEM-4 CPNE function have opposite effects gap junction coupling. Statistical differences were determined as follows: Kruskal-Wallis test with Dunn’s multiple comparisons test (B); and one-way ANOVA with Tukey’s multiple comparisons test (C,D). Values that differ significantly are indicated (ns, not significant; **, *p* <0.01; ****, *p* <0.0001). Error bars indicate SEM. Sample sizes for each genotype are indicated in all figure panels.

To further address GEM-4’s role, we asked if constitutive GEM-4/CPNE expression in body muscles increases excitability, thereby mimicking the *mef-2* mutant phenotype. For this analysis, we used a single copy transgene (*nuSi697)* containing a muscle specific promoter (*pat-10*) driving expression of a *gem-4* cDNA, hereafter designated *gem-4(muscle excess, mxs)*. Consistent with this idea, the *gem-4(mxs)* transgene increased the evoked AP rate (Fig. 6A,C,D), depolarized the RMP (Fig. 6E), decreased C_m_ (Fig. 6F), increased R_in_ (Fig. 6B,G), and decreased the gap junction coupling coefficient (Fig. 7A-D). By contrast, *gem-4(mxs)* had no effect on SHK-1 current (Fig. 6, Supplement 1A,C). Thus, decreasing and increasing GEM-4 function produced reciprocal effects on muscle excitability, passive membrane properties, and gap junction coupling. The effects of *gem-4(mxs)* on AP rate (Fig. 6, Supplement 1D,E), RMP (Fig. 6, Supplement 1F), C_m_ (Fig. 6, Supplement 1G), and R_in_ (Fig. 6, Supplement 1H) were eliminated in *unc-9* mutants, suggesting that GEM-4 controls excitability and passive membrane properties via changes in gap junction coupling. Collectively, these results strongly support the idea that GEM-4 mediates the effects of MEF-2 and CRH-1/CREB on gap junction coupling.

An alternative explanation for these findings is that GEM-4/CPNE regulates CRH-1/CREB mediated transcription, thereby explaining the GEM-4’s effects on muscle excitability. We did several experiments to address this possibility. First, we found that *gem-4(mxs)* did not alter expression of the *gem-4(nu858* mSG::H2B) reporter (Fig. 6, Supplement 2A-B). Second, the effects of *gem-4(mxs)* on AP rate (Fig. 6, Supplement 2C-D), RMP (Fig. 6, Supplement 2E), C_m_ (Fig. 6, Supplement 2F), and R_in_ (Fig. 6, Supplement 2G) were not eliminated in *crh-1* CREB mutants. Third, MEF-2’s effects on SHK-1 current were eliminated in *crh-1* CREB mutants (Fig. 1, Supplement 1G-I) but were unaffected by mutations decreasing or increasing GEM-4 activity (Fig. 6, Supplement 1A-C). Thus, changes in GEM-4 function impacted just a subset of *crh-1* mutant muscle phenotypes. Collectively, these results suggest that changes in CRH-1 transcriptional activity are unlikely to account for GEM-4’s effects on muscle excitability.

### MEF-2 and GEM-4 remodel gap junctions

GEM-4 CPNE could decrease gap junction coupling by a variety of mechanisms. GEM-4 could directly bind to innexins (e.g. UNC-9), inhibiting their function. Alternatively, GEM-4 could alter the number of gap junctions, or their subunit composition. To distinguish between these possibilities, we analyzed the distribution of endogenous gap junctions in the ventral nerve cord. Gap junctions were detected using the *unc-1(amz20* GFP) allele, which contains a GFP tag inserted at the N-terminus of the endogenous *unc-1* gene ^66^. UNC-1 Stomatin binds directly to UNC-9 and is required for gap junction coupling ^59,66,67^. UNC-1(amz20 GFP) exhibited robust punctate fluorescence in the ventral nerve cord, which was dramatically reduced in *unc-9* innexin mutants (Fig. 8A-D). These results demonstrate that UNC-1(amz20 GFP) localization can be used assess the distribution of endogenous UNC-9-UNC-1 complexes in the ventral cord. UNC-1 puncta density (puncta number/linear distance in the ventral cord) and puncta size were significantly reduced in *mef-2* mutants (Fig. 8A-C), indicating a decreased number of UNC-9-UNC-1 complexes. The decreased UNC-1 puncta density and size seen in *mef-2* mutants was eliminated in *mef-2;gem-4* double mutants (Fig. 8A-C), demonstrating that GEM-4 is required for MEF-2 regulation of UNC-9-UNC-1 complexes. Taken together, these results suggest that MEF-2 and GEM-4 regulate gap junction coupling of body muscles by controlling the number of UNC-9-UNC-1 complexes.

**Figure 8.**
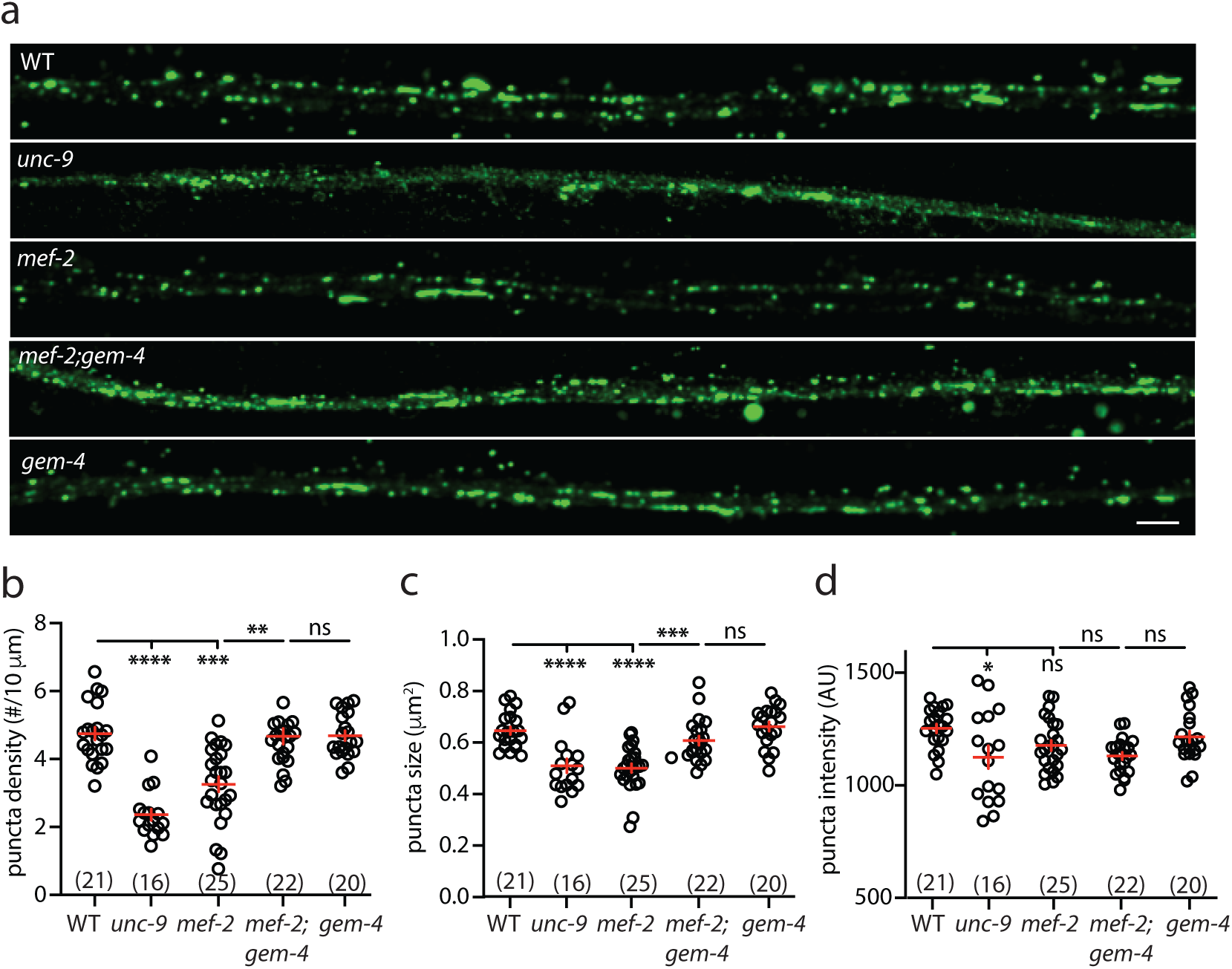
MEF-2 and GEM-4 remodel gap junction structure. UNC-1(amz20 GFP) Stomatin fluorescence was analyzed in the ventral nerve cord of adults of the indicated genotypes. Representative images (A), mean puncta density (B), mean puncta size (C), and mean puncta intensity (D) are shown. Representative images (A) and mean *gem-4* expression (B) are shown in the indicated genotypes. UNC-1 puncta density, size, and intensity were significantly decreased in *unc-9* mutants, indicating that UNC-1 localization can be used to analyze the distribution of UNC-9 containing gap junctions. UNC-1 puncta density and size were also significantly decreased in *mef-2* mutants, and these effects were eliminated in *mef-2;gem-4* double mutants. Statistical differences were determined by one-way ANOVA with Tukey’s multiple comparisons test. Values that differ significantly are indicated (ns, not significant; *, *p* <0.05; **, *p* <0.01; ***, *p* <0.001;****, *p* <0.0001). Error bars indicate SEM. Sample sizes for each genotype are indicated in all figure panels. Scale bar indicates 4 μm.

## Discussion

Although several identified ART targets promote synaptic plasticity, little is known about the transcriptional targets controlling intrinsic excitability. Here, we identify *gem-4* CPNE as an ART target that promotes plasticity of intrinsic excitability in *C. elegans* body muscles. These results show that ART regulation of excitability is ancient and occurs in both neurons and muscles.

Our results lead to five principal conclusions. First, MEF-2 decreases the intrinsic excitability of body muscles by repressing CRH-1 transcriptional targets. Second, MEF-2 inhibits muscle excitability by increasing gap junction coupling. Third, MEF-2 and CRH-1 reciprocally regulate expression of *gem-4* CPNE. Fourth, increased *gem-4* CPNE expression is both necessary and sufficient for the decreased gap junction coupling and increased muscle excitability seen in *mef-2* mutants. And fifth, GEM-4 inhibits gap junction coupling by structurally remodeling UNC-9 containing gap junctions. Below we discuss the significance of these findings.

### MEF-2 and CRH-1 antagonistically regulate excitability

Several prior studies suggest that ART regulates intrinsic excitability of neurons^36^. For example, CREB expression has been shown to increase excitability of several classes of neurons, in both mice and *Aplysia* ^37–41^. The transcriptional targets linking CREB expression to altered excitability were not identified in these studies. Here we describe a signaling pathway that functions in body muscles to link ART to changes in intrinsic excitability (Fig. 9A). We propose that increased muscle activity activates type 1 calcium and calmodulin (CaM) dependent protein kinase (CAMK1), which activates CRH-1 CREB ^8,9^ and inactivates MEF-2 mediated repression ^11^, thereby inducing *gem-4* CPNE expression.

**Figure 9.**
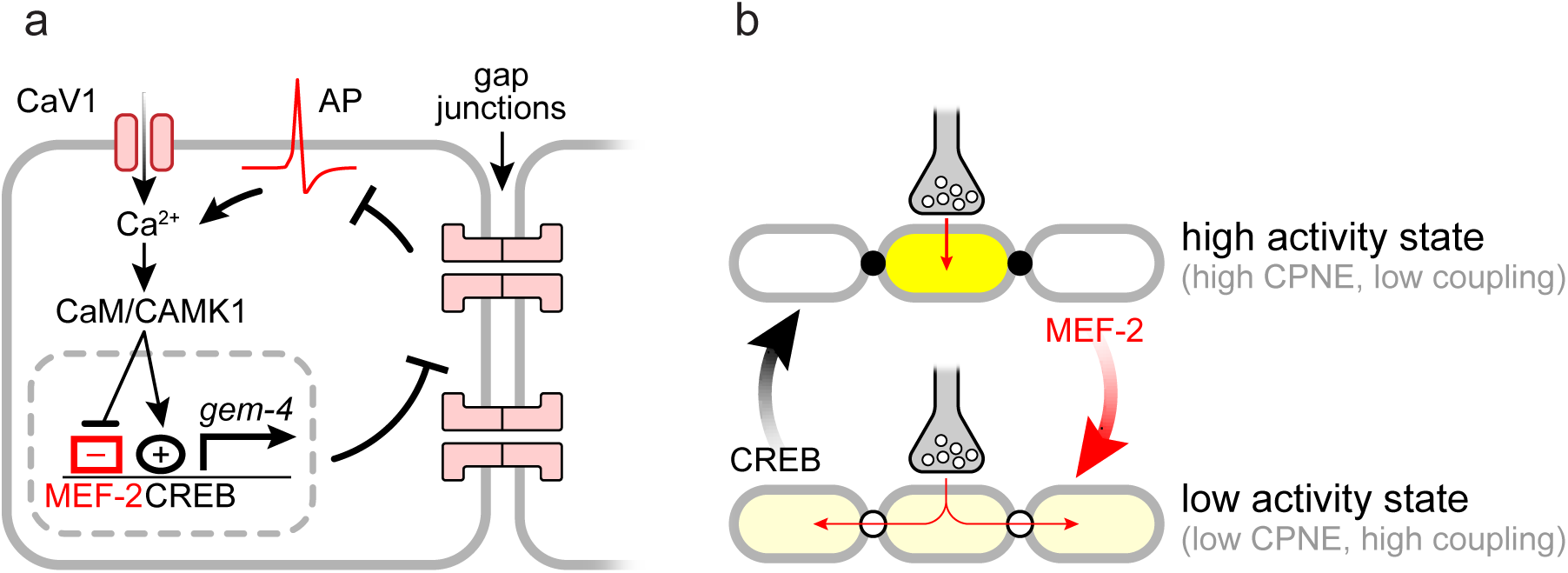
A model for GEM-4 regulation of muscle excitability. (A) Adjacent gap junctioned body muscles are shown. Following depolarization, calcium rises in the cytoplasm activating CMK-1 CAMK1. CMK-1 phosphorylation activates CRH-1 CREB ^8,9^ and disrupts MEF-2/HDAC4 ^11^ complexes, thereby inducing *gem-4* CPNE expression. GEM-4 CPNE expressed in muscles remodels gap junctions (decreasing the number of UNC-9-UNC-1 complexes), thereby increasing membrane resistance and intrinsic excitability. (B) A model illustrating how MEF-2 and CRH-1 promote inactive and active muscle states via activity induced CPNE expression. In active muscles, CRH-1 activity is high and MEF-2 repression is low, leading to high CPNE expression, decreased electrical coupling, and heightened excitability. In inactive muscles, CRH-1 activity is low and MEF-2 repression is high, leading to low CPNE expression, increased electrical coupling, and diminished excitability.

Increased GEM-4 levels in body muscles decreases gap junction coupling, thereby altering passive membrane properties (decreasing C_m_ and increasing R_in_) and increasing intrinsic excitability (increased AP rate) (Fig. 9A). Many results support this model. Inactivating MEF-2 increases muscle excitability and this effect is blocked in *mef-2;crh-1* double mutants, implying that MEF-2 inhibits excitability by repressing expression of CRH-1 CREB transcriptional targets. Muscle specific *mef-2* knockouts and muscle specific *crh-1* rescues indicate that MEF-2 and CRH-1 act in body muscles to control excitability and passive membrane properties. Increased excitability (and altered passive membrane properties) in *mef-2* mutants were eliminated by mutations inactivating *gem-4* CPNE, whose expression is repressed by MEF-2 and activated by CRH-1 CREB. Constitutive *gem-4* CPNE expression (in transgenic animals) mimics the *mef-2* mutant phenotypes of increased muscle excitability and altered passive membrane properties and these effects were not eliminated by mutations inactivating CRH-1 CREB, demonstrating that GEM-4 acts downstream of both MEF-2 and CRH-1. By contrast, the increased excitability resulting from *mef-2* inactivation or constitutive GEM-4 expression was prevented by mutations inactivating the *unc-9* innexin, indicating that UNC-9 containing gap junctions function downstream of MEF-2 and GEM-4. Gap junction coupling of body muscles is significantly reduced by *mef-2* inactivation (and is re-instated in *mef-2;gem-4* double mutants) and coupling is reduced by constitutive GEM-4 expression in muscles. Finally, constitutive GEM-4 expression (in *mef-2* mutants) decreases the number of UNC-9-UNC-1 complexes in the ventral nerve cord, which likely accounts for the decreased gap junction coupling because UNC-1 binding to UNC-9 is required for coupling ^59,67^. This decrease in UNC-9-UNC-1 complexes could result from decreased UNC-1 binding to UNC-9, or a decrease in expression, trafficking, or localization of UNC-1 or UNC-9 at muscle contact sites. Collectively, these results strongly support this model (Fig. 9A).

Because CBP, MEF2, and HDAC4 mutations are associated with several neurodevelopmental disorders^21–24^, it will be interesting to see if ART induced changes in intrinsic excitability contributes to the pathophysiology and comorbid phenotypes associated with these disorders.

### A new function for Copines

Copines are calcium binding proteins containing two C2 domains (C2A and C2B) and a VWA domain. Copines are widely conserved across eukaryotic phylogeny including protists, plants, and humans. Humans have 9 copine genes while *C. elegans* has 7 copines. Copines are proposed to play several roles in membrane trafficking, potentially via their interaction with the endosomal Rab protein RAB11 ^68^. Copines regulate synaptic vesicle exocytosis ^69^ and trafficking of several ion channels including TRP channels ^70,71^, synaptic GluA receptors ^63^, and nicotinic acetylcholine receptors ^72^. Copines also control the morphology of dendritic spines ^62,73^. Our results identify GEM-4/Copine as a physiologically important MEF2 and CREB target that regulates gap junctions. As proposed for other copines, GEM-4 could regulate excitability by altering the intracellular trafficking of critical proteins that regulate gap junctions. Because mouse CPNE6 expression is activity induced in the hippocampus ^61^, it will be interesting to see if this copine mediates ART’s effects of on hippocampal neuron excitability. Thus, copines have the capacity to control spine morphology, intrinsic excitability, and the trafficking of multiple cargo proteins (including synaptic GluA and ACh receptors) ^63,70–73^; consequently, ART induced copine expression could provide a flexible mechanism to promote long lasting structural, synaptic, and 18 intrinsic plasticity. Consistent with a role in late LTP, mouse CPNE6 is required to maintain potentiated transmission an hour after the inducing stimulus ^62^.

Prior studies showed that gap junction coupling is regulated by a variety of mechanisms^74,75^. In particular, cytoplasmic calcium can either activate or inhibit gap junction coupling ^76–80^. In some cases, these effects are mediated by calcium liganded calmodulin binding directly to connexin proteins ^81^. Gap junction coupling can also be regulated by phosphorylation or dephosphorylation of connexins ^82–84^. In *C. elegans*, UNC-1/Stomatin binding to UNC-9 innexins promote gap junction coupling ^67^. Our results identify GEM-4 Copine as a regulator of gap junction coupling, and highlight the potential importance of gap junction remodeling as a mechanism to regulate intrinsic excitability.

### Activity induced potentiation of muscle excitability

Decades of prior work has shown that many aspects of neuron function and development (including synapse development, synaptic strength, and intrinsic excitability) is regulated by preceding activity levels. Our results suggest that a similar mechanism exists in muscles whereby prior activity instructs changes in intrinsic excitability by controlling gap junction coupling (Fig. 9). It is interesting that hippocampal LTP and intrinsic plasticity of muscle excitability both require activity induced CPNE expression^62,63^, suggesting that these forms of neuron and muscle plasticity are mediated by a shared cell biological mechanism.

Use dependent plasticity of muscle excitability could have several important physiological functions. MEF2 and CREB could orchestrate switching between active and inactive muscle states (Fig. 9B). In this scenario, inactive muscles would have low CRH-1/CREB activity, high MEF-2 repression activity, low *gem-4* CPNE expression, high gap junction coupling, and consequently diminished excitability. By contrast, active muscles have high CRH-1/CREB activity, low MEF-2 repression activity, high *gem-4* expression, and exhibit the converse effects on gap junction coupling and excitability.

This mechanism could also alter the development of neuromuscular junctions (NMJs). NMJ development is often regulated by prior activity, whereby increased activity promotes growth of NMJ structure ^85,86^. The mechanism we describe here could enhance NMJ growth in active muscles (by increasing muscle excitability) and could make NMJ growth input selective (by decreasing gap junction coupling between muscles). Further experiments will be required to test these and other ideas for how use dependent muscle excitability impacts the function and development of motor circuits.

## Supporting information

Supplemental Data Set

## Acknowledgements

We thank the following for strains, advice, reagents, and comments on the manuscript: *C. elegans* genetics stock center (CGC), S. Mitani, Jeremy Dittman, Nathan Harris, Sam Bates, Piali Sengupta, and members of the Kaplan lab. We especially thank L. Orefice for providing the multiclamp amplifier used for the paired muscle recordings, A. Bhattacharya for providing the *unc-1(amz20* GFP) strain, and J. Dittman for providing the model diagram shown in Figure 9. This work was supported by NIH research grants to J.K. (NS32196). The CGC is funded by the NIH Office of Research Infrastructure Programs (P40 OD010440).

## Author Contributions

L.G. did all experiments, and constructed strains. E.P. collected preliminary data indicating increased muscle excitability in *mef-2* mutants. S.N. designed and isolated CRISPR alleles and constructed strains. J.M.K. supervised and acquired funding. J.M.K., L.G., and S.N. wrote and revised the paper.

## Experimental Methods

### Strains

Strain maintenance and genetic manipulation were performed as described ^87^. Animals were cultivated at room temperature (∼22°C) on agar nematode growth media seeded with OP50 bacteria. All strains used are listed in Table 1. Transgenic animals were prepared by microinjection. Single copy transgenes were isolated by the MoSCI and miniMoS techniques ^88,89^.

**Table 1.**
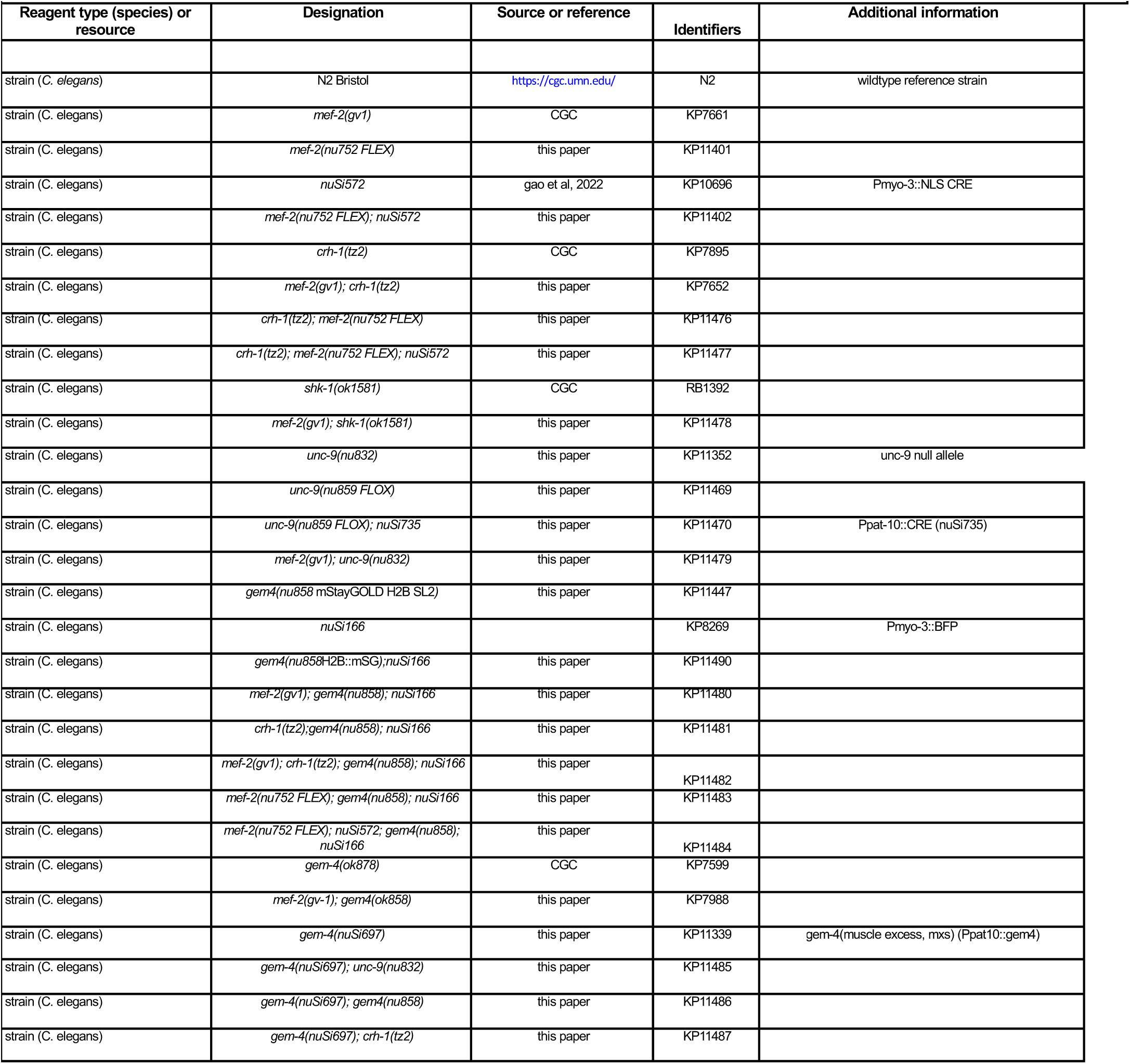
Strains used in this study.

| Reagent type (species) or resource | Designation | Source or reference | Identifiers | Additional information |
| --- | --- | --- | --- | --- |
| strain (*C. elegans*) | N2 Bristol | https://cgc.umn.edu/ | N2 | wildtype reference strain |
| strain (*C. elegans*) | *mef-2(gv1)* | CGC | KP7661 |  |
| strain (*C. elegans*) | *mef-2(nu752 FLEX)* | this paper | KP11401 |  |
| strain (*C. elegans*) | *nuSi572* | gao et al, 2022 | KP10696 | Pmyo-3::NLS CRE |
| strain (*C. elegans*) | *mef-2(nu752 FLEX); nuSi572* | this paper | KP11402 |  |
| strain (*C. elegans*) | *crh-1(tz2)* | CGC | KP7895 |  |
| strain (*C. elegans*) | *mef-2(gv1); crh-1(tz2)* | this paper | KP7652 |  |
| strain (*C. elegans*) | *crh-1(tz2); mef-2(nu752 FLEX)* | this paper | KP11476 |  |
| strain (*C. elegans*) | *crh-1(tz2); mef-2(nu752 FLEX); nuSi572* | this paper | KP11477 |  |
| strain (*C. elegans*) | *shk-1(ok1581)* | CGC | RB1392 |  |
| strain (*C. elegans*) | *mef-2(gv1); shk-1(ok1581)* | this paper | KP11478 |  |
| strain (*C. elegans*) | *unc-9(nu832)* | this paper | KP11352 | unc-9 null allele |
| strain (*C. elegans*) | *unc-9(nu859 FLOX)* | this paper | KP11469 |  |
| strain (*C. elegans*) | *unc-9(nu859 FLOX); nuSi735* | this paper | KP11470 | Ppat-10::CRE (nuSi735) |
| strain (*C. elegans*) | *mef-2(gv1); unc-9(nu832)* | this paper | KP11479 |  |
| strain (*C. elegans*) | *gem4(nu858 mStayGOLD H2B SL2)* | this paper | KP11447 |  |
| strain (*C. elegans*) | *nuSi166* |  | KP8269 | Pmyo-3::BFP |
| strain (*C. elegans*) | *gem4(nu858H2B::mSG); nuSi166* | this paper | KP11490 |  |
| strain (*C. elegans*) | *mef-2(gv1); gem4(nu858); nuSi166* | this paper | KP11480 |  |
| strain (*C. elegans*) | *crh-1(tz2); gem4(nu858); nuSi166* | this paper | KP11481 |  |
| strain (*C. elegans*) | *mef-2(gv1); crh-1(tz2); gem4(nu858); nuSi166* | this paper | KP11482 |  |
| strain (*C. elegans*) | *mef-2(nu752 FLEX); gem4(nu858); nuSi166* | this paper | KP11483 |  |
| strain (*C. elegans*) | *mef-2(nu752 FLEX); nuSi572; gem4(nu858); nuSi166* | this paper | KP11484 |  |
| strain (*C. elegans*) | *gem-4(ok878)* | CGC | KP7599 |  |
| strain (*C. elegans*) | *mef-2(gv-1); gem4(ok858)* | this paper | KP7988 |  |
| strain (*C. elegans*) | *gem-4(nuSi697)* | this paper | KP11339 | gem-4(muscle excess, mxs) (Ppat10::gem4) |
| strain (*C. elegans*) | *gem-4(nuSi697); unc-9(nu832)* | this paper | KP11485 |  |
| strain (*C. elegans*) | *gem-4(nuSi697); gem4(nu858)* | this paper | KP11486 |  |
| strain (*C. elegans*) | *gem-4(nuSi697); crh-1(tz2)* | this paper | KP11487 |  |

### CRISPR alleles

CRISPR alleles were isolated as described ^90^. Briefly, Cas9 protein and guide RNAs were ordered from IDT. Repair templates shorter than 200bp consisted of ss ULTRAMER oligos from IDT. Longer repair templates were PCR amplified from a plasmid and melted before adding to the injection mix. The injection mix included the pRF4 *rol-6*(gf) plasmid. Injected animals were singled and 96 F1 progeny were singled from plates that contained rollers, allowed to starve out the plate, and were then screened by PCR for the expected change.

Muscle specific knockouts were performed using CRE inactivated alleles. CRE recombinase was expressed in muscles (using either the *myo-3* or *pat-10* promoter). Muscle specific *mef-2* knockouts were performed by introducing a stop cassette into the first intron of the endogenous locus in the ON configuration (i.e. in the opposite orientation of the *mef-2* gene), creating the *mef-2(nu752, flex ON)* allele. The stop cassette consists of a synthetic exon (containing a consensus splice acceptor sequence and stop codons in all reading frames) followed by a 3’ UTR and transcriptional terminator, as described ^91^. The stop cassette is flanked by FLEX sites (which are modified loxP sites that mediate CRE induced inversions) ^92^. In this manner, *mef-2* expression in body muscles is reduced following CRE expression with the *myo-3* promoter (*nuSi572*). For *unc-9*, LoxP sites flanking the first exon were inserted into the endogenous locus, hereafter *unc-9(nu859* Flox), and CRE was expressed in muscles (using the *pat-10* promoter, *nuSi735*).

To construct a *gem-4* transcriptional reporter, the endogenous *gem-4* locus was converted into an operon containing an upstream cistron expressing histone H2B fused to the green fluorescent protein mStayGold (mSG), creating the *gem-4(nu858* mSG::H2B) allele. This allele was used to quantify *gem-4* expression in body muscles. Transgenes expressing nuclear localized fluorescent proteins were used to identify body muscle nuclei (*nuSi166*, NLS-BFP, *myo-3* promoter). Expression of *gem-4* was assessed by measuring the mSG/BFP fluorescence ratio in individual muscle nuclei.

The *unc-9(nu832)* null allele is a 3825 nucleotide deletion spanning all exons of the *unc-9* locus.

### Fluorescence imaging

Worms were immobilized on 10% agarose pads with 3 µl of 0.1 µm diameter polystyrene microspheres (Polysciences 00876-15, 2.5% w/v suspension). Images were taken with a Nikon A1R confocal, using a 60X/1.49 NA oil objective. For *gem-4(nu858* mSG::H2B) imaging, image volumes spanning muscle nuclei were collected (∼20 planes/volume, 0.225 μm between planes, and 0.29 μm /pixel). Maximum intensity projections for each volume were auto-thresholded, and nuclei were identified as round fluorescent objects, using analysis of particles. Mean fluorescent intensity in each nucleus was analyzed in the raw images. For *unc-1(amz20* GFP) imaging, image volumes spanning the ventral nerve cord were collected (∼20 planes/volume, 0.225 μm between planes, and 0.144 μm /pixel) and deconvoluted using Nikon 3D Deconvolution. Maximum intensity projections of deconvoluted images for each volume were auto-thresholded, and puncta were identified as round fluorescent objects (area > 0.2 μm^2^), using analysis of particles. Mean puncta density (puncta number/ventral cord length), size, and fluorescent intensity was analyzed. All image analysis was done using FIJI.

### Electrophysiology

Whole-cell patch-clamp measurements were performed using a Axopatch 200B amplifier with pClamp 10 software (Molecular Devices). For paired muscle recordings, adjacent ventral body muscles were subjected to whole-cell patch clamp using a MultiClamp 700B amplifier with MultiClamp 700B commander and pClamp 10 software (Molecular Devices). The data were sampled at 10 kHz and filtered at 5 kHz. All recordings were performed at room temperature (∼19-21 °C).

#### Muscle AP recordings

The bath solution contained (in mM): NaCl 140, KCl 5, CaCl_2_ 5, MgCl_2_ 5, dextrose 11 and HEPES 5 (pH 7.2, 320 mOsm). The pipette solution contained (in mM): Kgluconate 120, KOH 20, Tris 5, CaCl_2_ 0.25, MgCl_2_ 4, sucrose 36, EGTA 5, and Na_2_ATP 4 (pH 7.2, 323 mOsm). Evoked APs were recorded in current clamp following -8pA to 10pA with 2pA step current injections. Cell resistance (R_in_) was measured using a -8 pA pulse injection. Cell capacitance (C_m_) was measured as follows: in voltage-clamp, a -30mV (-60mV to -90mV) voltage-step is applied to the cell, and the resulting current is measured. Cell capacitance was calculated as C_m_=Q/ΔV, where Q is the total transient membrane charge (measured using pClamp 10) and ΔV is the voltage applied to the cell (-30mV).

#### Muscle K^+^ current recordings

The bath solution contained (in mM): NaCl 140, KCl 5, CaCl_2_ 5, MgCl_2_ 5, dextrose 11 and HEPES 5 (pH 7.2, 320 mOsm). For SHK-1 recordings, the pipette solution contained (in mM): Kgluconate 120, KOH 20, Tris 5, CaCl_2_ 0.25, MgCl_2_ 4, sucrose 36, EGTA 5 and Na2ATP 4 (pH 7.2, 323 mOsm). For SLO-2 recordings, the pipette solution contained (in mM): KCl 120, KOH 20, Tris 5, CaCl_2_ 0.25, MgCl_2_ 4, sucrose 36, EGTA 5, and Na_2_ATP 4 (pH 7.2, 323 mOsm). SLO-2 current was isolated by recording in *shk-1* mutants. The voltage-clamp protocol consisted of -60mV for 50ms, -90mV for 50 ms, test voltage (from -60mV to +60mV) 200 ms. In figures, we show outward currents evoked at +30 mV, which corresponds to the peak amplitude of muscle APs.

#### Muscle CaV recordings

The bath solution contained (in mM): TEA-Cl 140, CaCl_2_ 5, MgCl_2_ 1, 4AP 3, glucose 10, sucrose 5 and HEPES 15 (pH 7.4, 330 mOsm). The pipette solution contained (in mM): CsCl 140, TEA-Cl 10, MgCl_2_ 5, EGTA 5 and HEPES 5 (pH 7.2, 320 mOsm). The voltage-clamp protocol consisted of -60mV for 50ms, -90mV for 50 ms, test voltage (from -60mV to +60mV) for 200 ms.

#### Paired muscle recordings

Adjacent body muscles L1L2 or R1R2 were subjected to whole cell patch clamp (as illustrated in Fig. 4g). V_m_ was measured in both the clamped (cell 1) and unclamped (cell 2) muscles in current clamp following -10, -20, and -30 pA injections into cell 1 for 2s.

### Statistical methods

Data graphing and statistics were performed in GraphPad Prism 11. No statistical method was used to select sample sizes. For normally distributed data, significant differences were assessed with unpaired t tests (for 2 groups) or one way ANOVA with Tukey’s multiple comparisons test (for >2 groups). For non-normal data, differences were assessed by Mann-Whitney (2 groups) or Kruskal-Wallis test with post-hoc Dunn’s multiple comparisons test (>2 groups). Specific statistical tests used are indicated in each figure legend. Data shown in each figure represent contemporaneous measurements from mutant and controls over a period of 1-2 weeks. For electrophysiology, data points represent mean values for muscle recordings (which were considered biological replicates). For imaging studies, data points represent mean puncta fluorescence values in individual animals (which were considered biological replicates). All data obtained in each experiment were analyzed, without any exclusions.

---

**Figure 1-Supplement 1.**
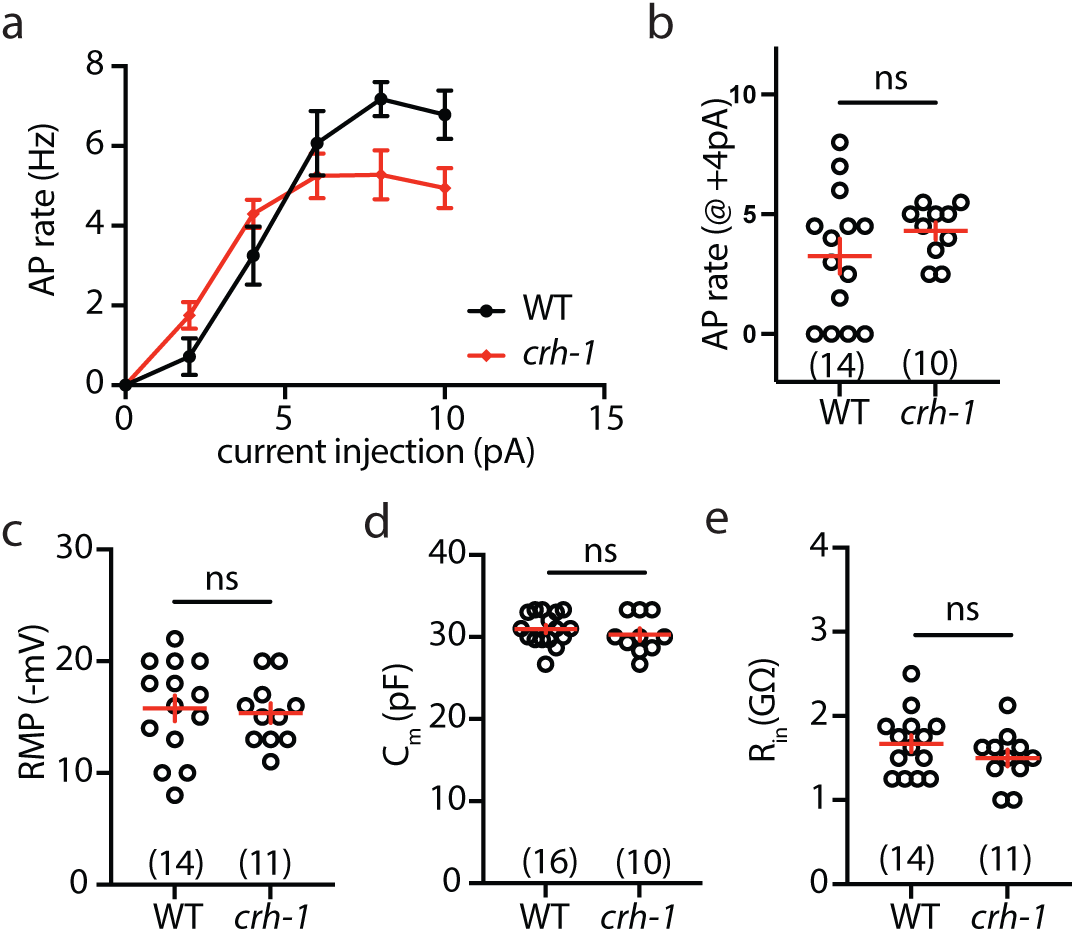
Muscle excitability and passive membrane properties are unaltered in *crh-1* CREB single mutants. Muscle excitability is compared in WT and *crh-1* CREB single mutants. Mean AP rate as a function of injected current (A), mean AP rate evoked by a +4 pA current (B), mean RMP (C), mean C_m_ (D), and mean R_in_ (E) are shown. No significant differences were observed by unpaired t test with Welch’s correction (ns, not significant). These results suggest that inactivating CRH-1/CREB alone was not sufficient to alter muscle excitability. Error bars indicate SEM. Sample sizes for each genotype are indicated in all figure panels.

**Figure 2-Supplement 1.**
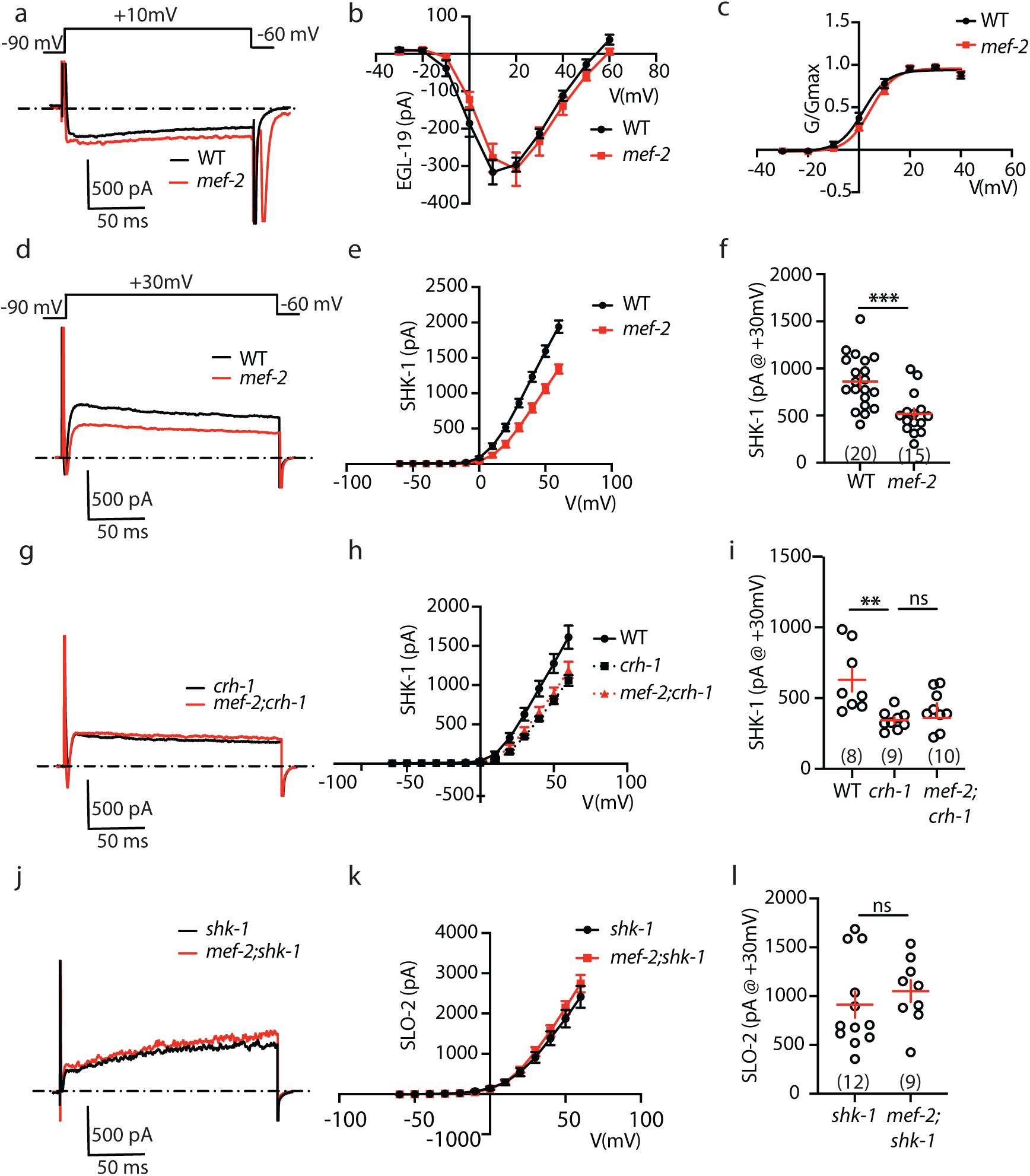
Analysis of voltage-activated currents in *mef-2* mutants. EGL-19 CaV1 (A-C), SHK-1 KCNA (D-I), and SLO-2 BK (J-L) currents were analyzed in *mef-2* null mutants. Representative traces (A,D,G,J), current amplitude as a function of membrane potential (B,E,H,K), conductance as a function of membrane potential (C), and current amplitude at +30 mV (F,I,L) are shown for the indicated genotypes. EGL-19 CaV1 current amplitude (A,B) and the voltage-dependence of EGL-19 activation (C) were unaffected in *mef-2* null mutants. SHK-1 KCNA current was significantly decreased in both *mef-2* (D-F) and *crh-1* (G-I) single mutants, and additive effects were not observed in *mef-2;crh-1* double mutants. Lack of additive effects in the double mutants suggest that MEF-2’s impact on SHK-1 channels is mediated by increased expression of one or more CRH-1 transcriptional target. SLO-2 BK current was unaffected in *mef-2* null mutants (J-L). Statistical differences were determined as follows: one-way ANOVA with Tukey’s multiple comparisons test (I); unpaired t test with Welch’s correction (F and L). Values that differ significantly are indicated (ns, not significant; **, *p* <0.01; ***, *p* <0.001). Error bars indicate SEM. Sample sizes for each genotype are indicated in all figure panels.

**Figure 2-Supplement 2.**
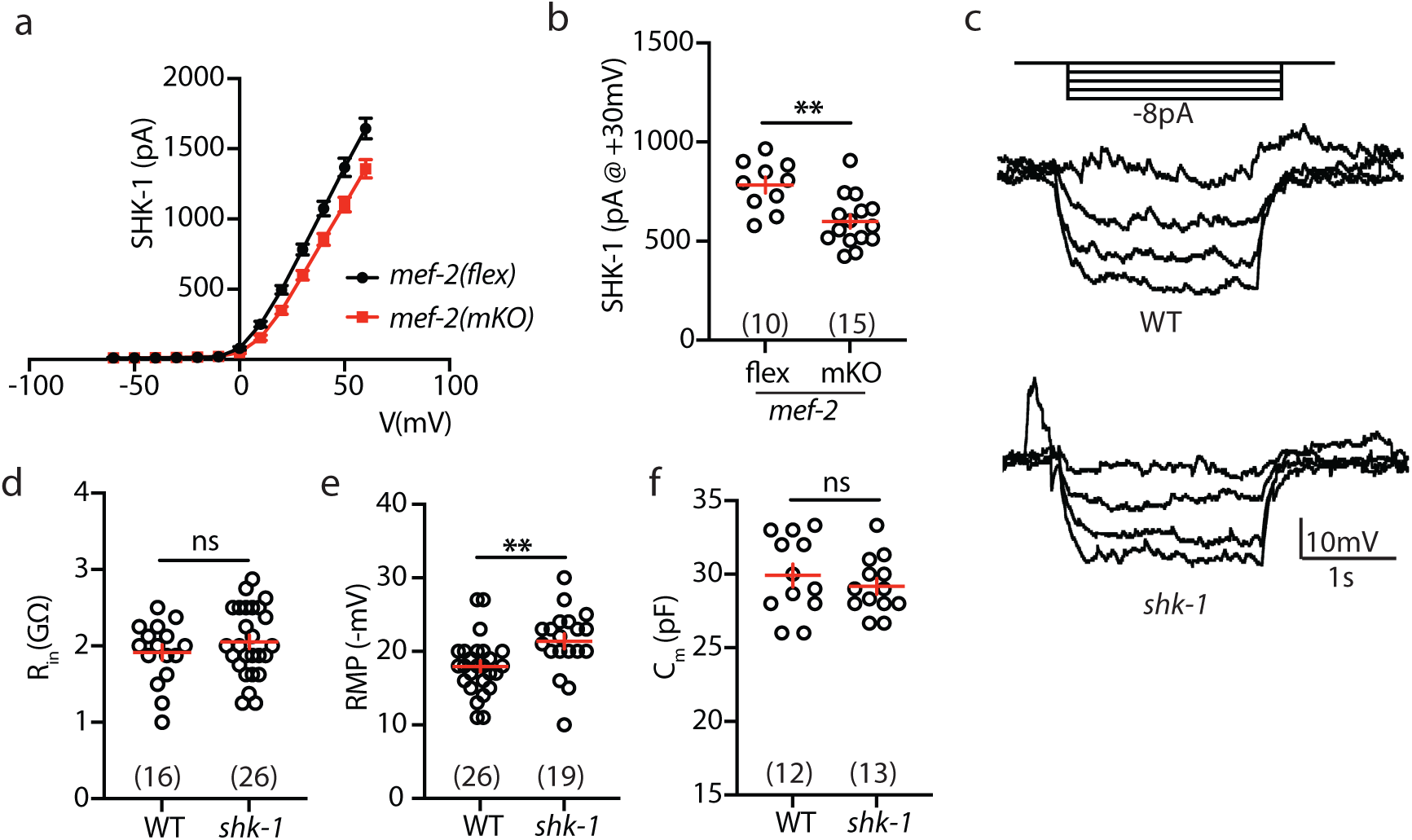
Decreased SHK-1 KCNA current is unlikely to account for the increased muscle excitability seen in *mef-2* mutants. (A-B) SHK-1 KCNA current is decreased in muscle specific *mef-2* knockouts (mKO), indicating that MEF-2 acts in body muscles to regulate SHK-1 KCNA channels. *mef-2(flex)* is the WT control (lacking CRE expression) for *mef-2(mKO)*. SHK-1 KCNA current as a function of membrane potential (A) and peak SHK-1 current amplitude evoked at +30 mV (B) are shown in the indicated genotypes. (C-F) Intrinsic muscle excitability was not altered in *shk-1* KCNA mutants. Representative traces of membrane potential changes evoked by hyperpolarizing current injections (C), mean input resistance (R_in_) (D), mean RMP (E), and mean cell capacitance (C_m_) (F) are compared in WT and *shk-1* null mutants. Statistical differences were determined by unpaired t test with Welch’s correction. Values that differ significantly are indicated (ns, not significant; **, *p* <0.01). Error bars indicate SEM. Sample sizes for each genotype are indicated in all figure panels.

**Figure 3-Supplement 1.**
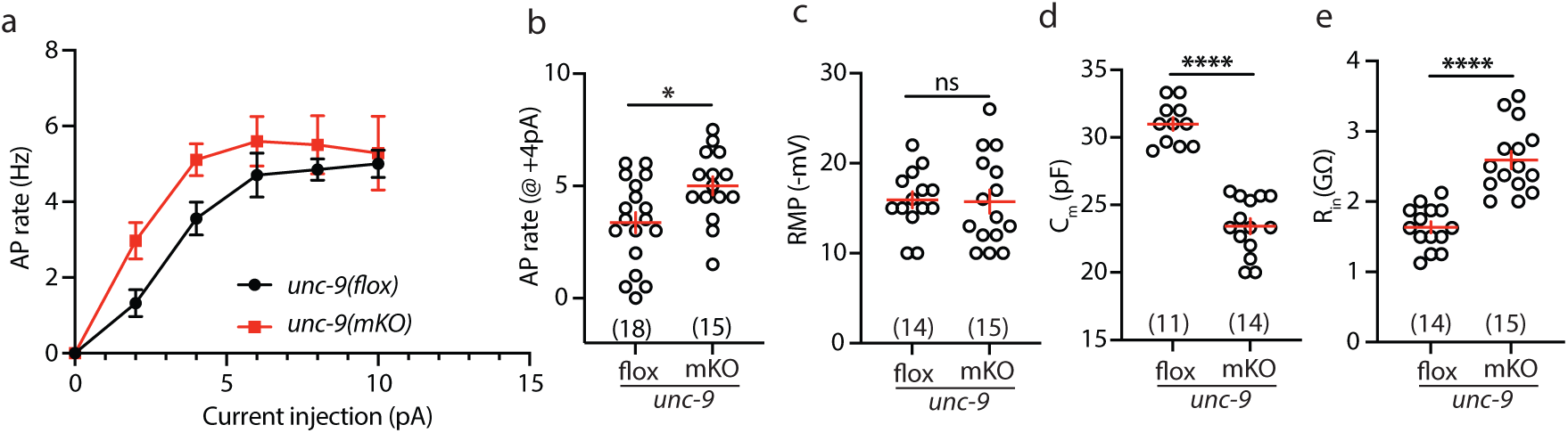
UNC-9 acts in body muscles to regulate intrinsic excitability. Muscle excitability is compared in *unc-9(flox)* and *unc-9(mKO)* mutants. *unc-9(flox)* is the WT control (lacking CRE expression) for *unc-9(mKO)*. Mean AP rate as a function of injected current (A), mean AP rate evoked by a +4 pA current (B), mean RMP (C), mean C_m_ (D), and mean R_in_ (E) are shown. Following CRE expression in body muscles, *unc-9(mKO)* mutants exhibited an increased evoked AP rate (A,B), a decreased C_m_ (D), and an increased R_in_ (E). These results suggest that inactivating *unc-9* selectively in body muscles was sufficient to increase excitability and to alter passive membrane properties. Statistical differences were determined by unpaired t test with Welch’s correction. Values that differ significantly are indicated (ns, not significant; *, *p* <0.05; ****, *p* <0.0001). Error bars indicate SEM. Sample sizes for each genotype are indicated in all figure panels.

**Figure 6-Supplement 1.**
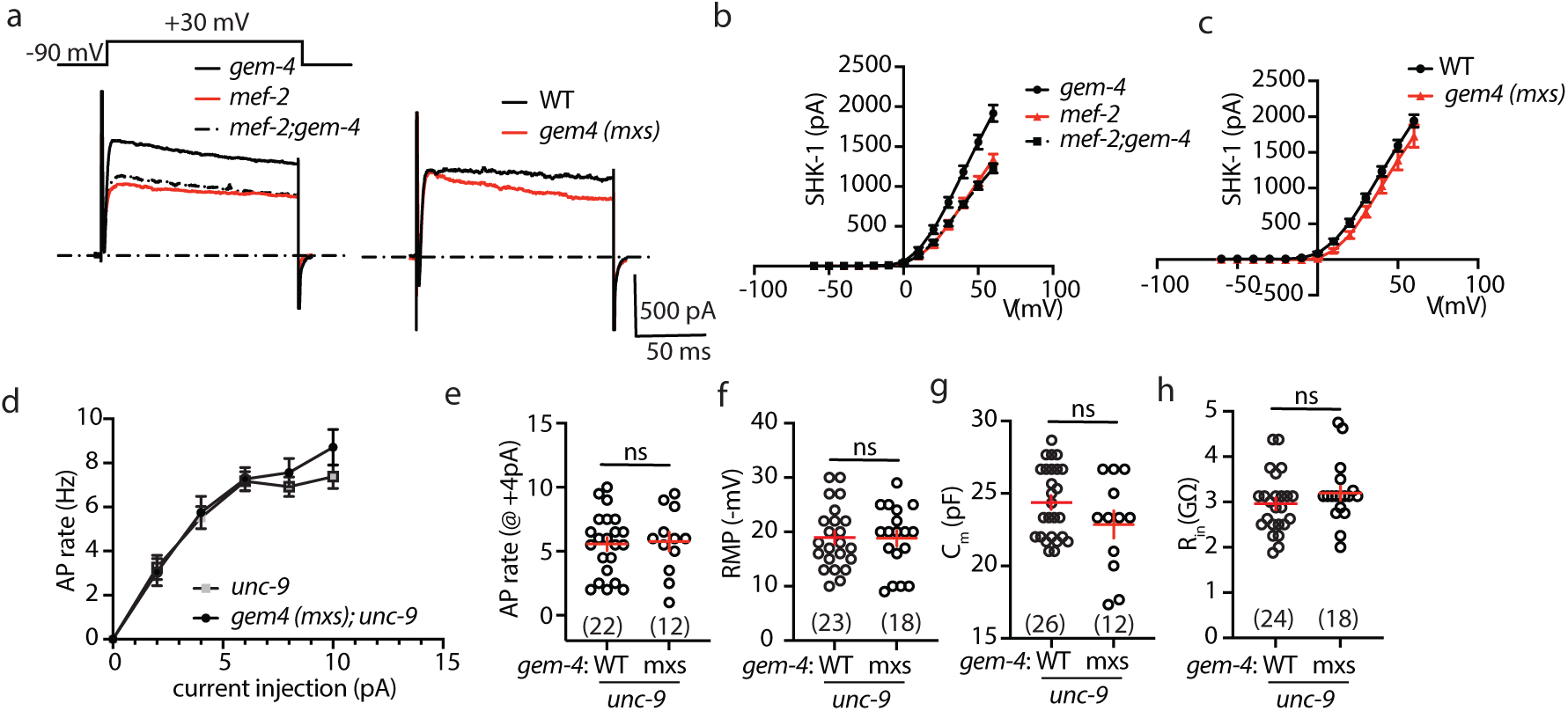
UNC-9 is required for GEM-4 CPNE’s effects on intrinsic excitability. (A-C) Increased and decreased GEM-4 CPNE function have no effect on SHK-1 KCNA current. Representative traces of SHK-1 current evoked at +30 mV (A) and SHK-1 current as a function of membrane potential (B,C) are shown in the indicated genotypes. (D-H) Increased GEM-4 CPNE expression has no effect on muscle excitability in *gem-4(mxs);unc-9* double mutants. These results suggest that changes in GAP junction coupling are required for GEM-4’s effects on excitability. Mean AP rate as a function of injected current (D), mean AP rate evoked by a +4 pA current (E), mean RMP (F), mean C_m_ (G), and mean R_in_ (H) are shown. No significant differences were observed by unpaired t test with Welch’s correction (ns, not significant). Error bars indicate SEM. Sample sizes for each genotype are indicated in all figure panels.

**Figure 6-Supplement 2.**
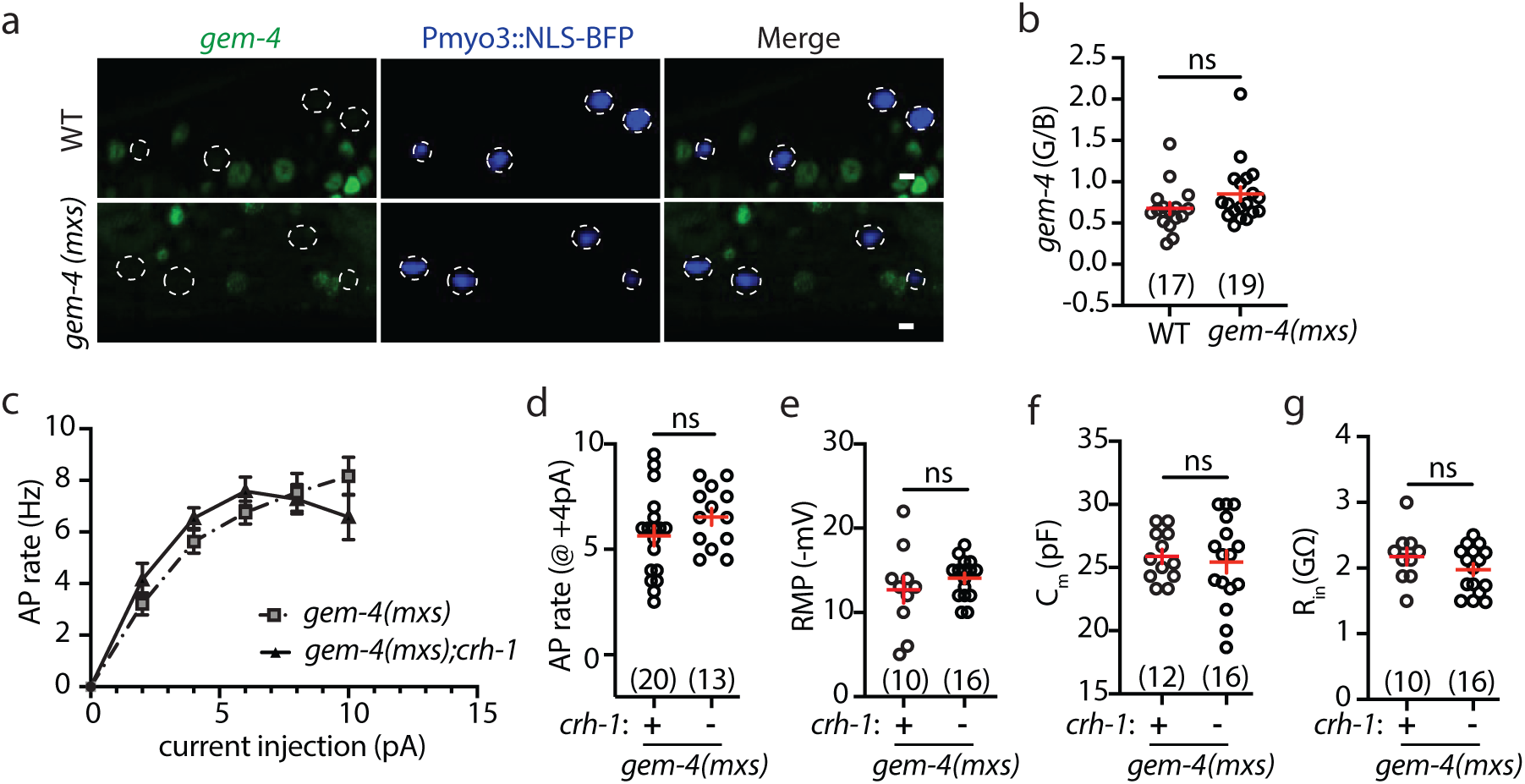
CRH-1 CREB is not required for GEM-4 CPNE’s effects on intrinsic excitability. (A-B) Increased GEM-4 CPNE had no effect on *gem-4(mSG::H2B)* expression. Representative images (A) and mean *gem-4* expression (B) are shown in the indicated genotypes. Body muscle nuclei (dashed circles) were identified by a transgene expressing nuclear localized BFP (NLS-BFP) with the *myo-3* promoter. (C-G) Increased GEM-4 CPNE’s effects on intrinsic muscle excitability and passive membrane properties were unaltered in *crh-1* CREB mutants. Mean AP rate as a function of injected current (C), mean AP rate evoked by +4 pA (D), mean RMP (E), mean C_m_ (F), and mean R_in_ (G) are shown. No significant differences were observed by unpaired t test with Welch’s correction (ns, not significant). Error bars indicate SEM. Sample sizes for each genotype are indicated in all figure panels. Scale bar indicates 4 μm.

